# Evolutionary origins and innovations sculpting the mammalian PRPS enzyme complex

**DOI:** 10.1101/2024.10.01.616059

**Authors:** Bibek R. Karki, Austin C. Macmillan, Sara Vicente-Muñoz, Kenneth D. Greis, Lindsey E. Romick, J. Tom Cunningham

## Abstract

The phosphoribosyl pyrophosphate synthetase (PRPS) enzyme conducts a chokepoint reaction connecting central carbon metabolism and nucleotide production pathways, making it essential for life^1,2^. Here, we show that the presence of multiple PRPS-encoding genes is a hallmark trait of eukaryotes, and we trace the evolutionary origins and define the individual functions of each of the five mammalian PRPS homologs – three isozymes (one testis-restricted)^3,4^ and two non-enzymatic associated proteins (APs)^5,6^ – which we demonstrate operate together as a large molecular weight complex capable of attaining a heterogeneous array of functional multimeric configurations. Employing a repertoire of isogenic fibroblast clones in all viable individual or combinatorial assembly states, we define preferential interactions between subunits, and we show that cells lacking PRPS2, PRPSAP1, and PRPSAP2 render PRPS1 into aberrant homo-oligomeric assemblies with diminished metabolic flux and impaired proliferative capacity. We demonstrate how numerous evolutionary innovations in the duplicated genes have created specialized roles for individual complex members and identify translational control mechanisms that enable fine-tuned regulation of PRPS assembly and activity, which provide clues into the positive and negative selective pressures that facilitate metabolic flexibility and tissue specialization in advanced lifeforms. Collectively, our study demonstrates how evolution has transformed a single PRPS gene into a multimeric complex endowed with functional and regulatory features that govern cellular biochemistry.

## Introduction

Phosphoribosyl pyrophosphate synthetase (PRPS) is an enzyme conserved across all forms of life, tracing back to the last universal common ancestor (LUCA)^7^. PRPS catalyzes the rate-limiting step in converting ribose-5-phosphate (R5P) to phosphoribosyl pyrophosphate (PRPP), a crucial precursor in the biosynthesis of nucleotides, amino acids, and lipids^1,2^. In mammals, there are three PRPS isoforms, which form a heteromeric complex with two additional homologs—PRPS-associated protein 1 (PRPSAP1) and PRPS-associated protein 2 (PRPSAP2)^3–6,8–10^. However, their evolutionary origins, the functional significance of these heteromeric assemblies and the specific roles of each homolog are still unclear. Here, we perform evolutionary analyses to investigate the origins of mammalian PRPS homologs and employ biochemical and genetic approaches to establish the structure-function relationships that influence assembly and activity of the mammalian PRPS complex.

Our findings trace the origins of mammalian PRPSAP2 to a gene duplication of PRPS1 in early Opisthokonta, and we identify a second gene duplication event in the ancestor of jawed vertebrates that produced PRPS2 and PRPSAP1, from PRPS1 and PRPSAP2, respectively. We demonstrate that the heteromeric PRPS enzyme complex is among the largest assemblies in mammalian cells, with different tissues achieving unique architectures by varying the stoichiometric expression of individual components. We show the critical importance of proper PRPS complex assembly for enzyme functionality within cells as improper assembly perturbs global metabolic flux and decreases cellular fitness. Additionally, we utilize genetic engineering to define the organization and assembly of the complex, establish preferential interactions among its members, and uncover regulatory mechanisms linking translational control to complex assembly. Our thorough phylogenomic analyses reveal multiple convergent evolutionary patterns giving rise to PRPSAP-like homologs throughout Amorphea, reinforcing the importance of PRPS enzymes operating as multimeric complexes in diverse eukaryotes.

## Results

### Evolutionary analysis of PRPS enzymes reveals gene duplication events and functional divergence in eukaryotes

Bacteria and Archaea species typically express a single PRPS enzyme^11–14^, whereas the few eukaryotic organisms studied to date possess multiple PRPS homologs^3,9,15–17^. We wondered whether this observation was extensible over the entire domain of eukaryotes. Sequence-based homology searches and genomic analyses, including more than 100 newly annotated sequences have confirmed that nearly all organisms in the eukaryotic domain contains multiple PRPS homologs (Fig.1a and Supplementary Table 1). Our extensive catalog of the essential enzyme PRPS can help reconstruct organismal phylogeny across different eukaryotic species and illustrate how evolutionary pressures contribute to the emergence of new properties over evolutionary timescales. We hypothesized that this increased repertoire of PRPS homologs imbues eukaryotes with enhanced metabolic adaptability by virtue of additional regulatability and increased biosynthetic capacity.

**Fig. 1:**
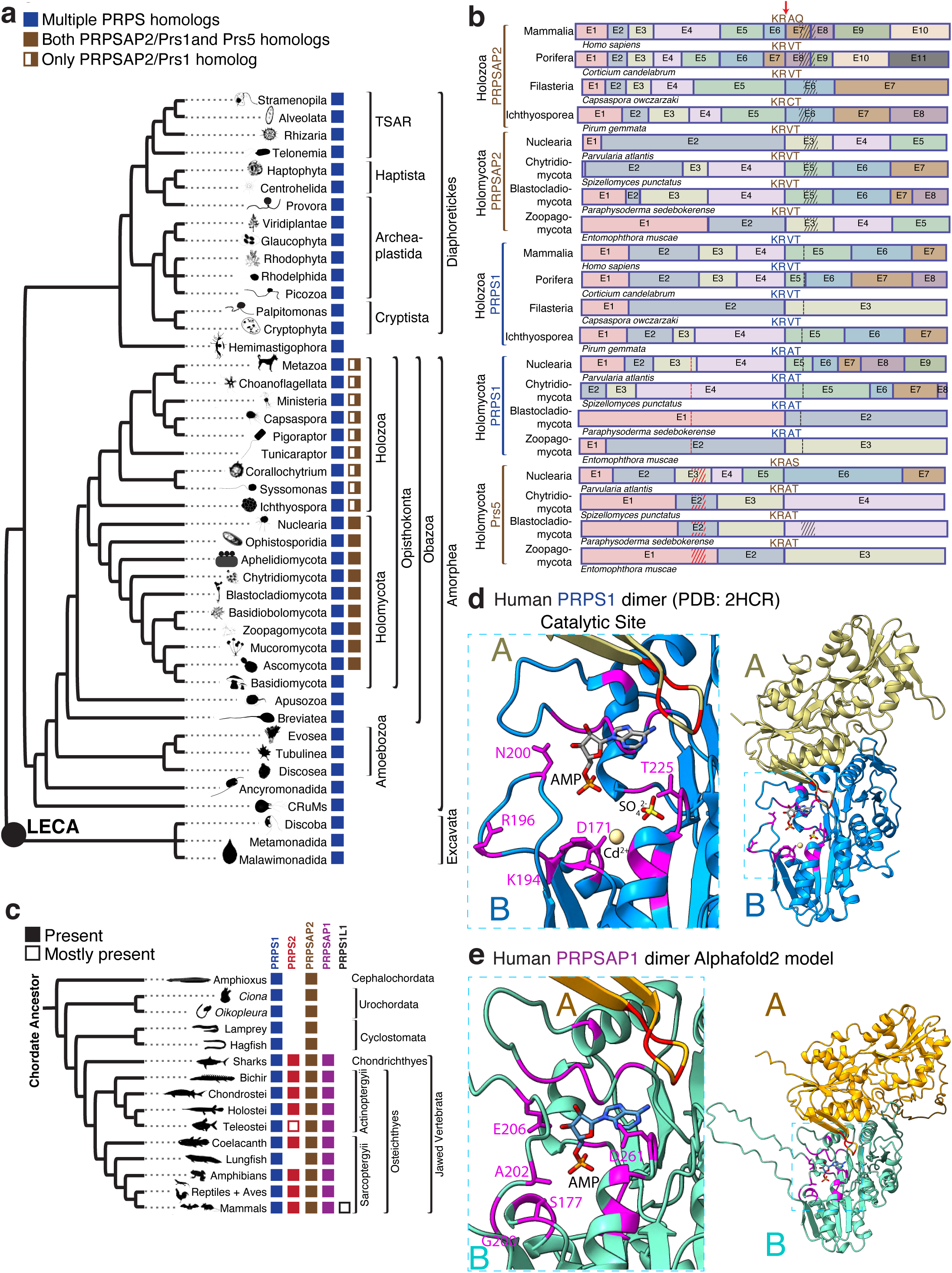
The encoding of multiple PRPS genes, including non-catalytic PRPSAPs, is a cardinal feature of eukaryotes. a) Phylogenetic distribution profiles of PRPS homologs in eukaryotes and PRPS homologs containing NHR in opisthokonts (presence/absence) are noted across the tree. PRPSAP2 represent orthologs of mammalian PRPSAP2 while Prs1 and Prs5 represent orthologs of *S. cerevisiae* Prs1 and Prs5, respectively. b) Analysis of conserved splice site junction among PRPS homologs across different representative organisms of Opisthokonta. Exons from each of the PRPS, PRPSAP2 and Prs5 encoding genes are merged to highlight the highly splice site junction with coding amino acids above. In PRPS1, red dots mark the regulatory flexible loop where insertions occur in Prs5 orthologs (red hatch marks), and black dots mark the catalytic flexible loop where insertions occur in PRPSAP2 orthologs (black hatch marks). Notably, some Prs5 orthologs possess additional insertions in the catalytic flexible loop. c) Phylogenetic distribution profiles of PRPS homologs in chordates (presence/absence) are noted across the tree. d) The structure of dimeric human PRPS1 (PDB ID: 2HCR), and a zoom in of the catalytic site highlighting metal binding site (Cd^2+^), AMP (representing the AMP moiety of the ATP), SO_4_ ^2-^ (representing the 5’-phosphate of R5P), and several conserved active site residues (magenta). D171 coordinates metal binding, K194 interacts with the ATP, R196 and T225 interact with the R5P, and N200 stabilizes the catalytic loop^22^. e) Predicted dimeric structure of human PRPSAP1 (Accession #AAH09012.1) from the AlphaFold2 model, and a zoom in highlights four non-conserved residues in PRPSAP1 (magenta) at the corresponding positions of residues for PRPS1 shown in (d). AMP was modeled into the dimer to denote the putative ATP binding site.

Given the well-established evolutionary trajectory from opisthokonts to mammals, supported by relatively complete molecular phylogenetic data, we used mammals, that harbor five PRPS homologs, as a model system to investigate the evolutionary origins and functional significance of these multiple homologs. In addition to the three isozymes– PRPS1, PRPS2, and PRPS1L1^3,4^ – mammals also possess PRPSAP1 and PRPSAP2^5,6^, which feature unique insertions in the catalytic flexible (CF) loop (referred to as non-homologous regions (NHRs)) compared to the sequences found in PRPS isozymes. Interestingly, this pattern of CF loop insertions is not exclusive to Holozoa but is also found in Holomycota (for example, *Saccharomyces cerevisiae* Prs1 and Prs5 homologs^17^) as well as some species from Amoebozoa, Apusozoa, and CRuMs (Supplementary Figs.1a and 1b). To determine whether the NHR-containing homologs in these Amorphean lineages originated from a common ancestor or represent cases of convergent evolution, we traced their evolutionary origins. Our analysis revealed evidence for independent origins through gene duplication events in Opisthokonta, Amoebozoa, Apusozoa, and CRuMs. PRPSAP2 orthologs (termed Prs1 in Holomycota) emerged in basal opisthokonts and share greater amino acid sequence homology with opisthokont PRPS1 than with PRPS proteins from non-opisthokont lineages (Supplementary Fig.1c). Notably, Prs5 orthologs, which contain insertions in the regulatory flexible loop, arose in the ancestor of Holomycota and are exclusive to this lineage (Fig.1a). Comparative gene structure analysis revealed that orthologs of opisthokont PRPS1, PRPSAP2 and Prs5 share a conserved splice site junction, despite over a billion years of divergent evolution, suggesting that a gene duplication event in the ancestral PRPS1 encoding gene led to the emergence of PRPSAP2 and Prs5 (Fig.1b). Similarly, PrsB orthologs from Evosea exhibit greater amino acid sequence homology and share a conserved splice site junction with Evosea PrsA (classical PRPS), supporting their independent origins and providing strong evidence of convergent evolution (Supplementary Figs.1d and1e). Here, we have uncovered one of the earliest genetic events that define Opisthokonta as well as distinguish Holomycota from Holozoa, thus showcasing the power of studying paralogous gene evolution.

Later in Holozoa evolution, another gene duplication occurred in the ancestor of jawed vertebrates, likely from a genome duplication event^18^, giving rise to PRPS2 and PRPSAP1 from PRPS1 and PRPSAP2, respectively (based on amino acid sequence homology summarized in Fig.1c). Comparative gene structure analysis of exon-intron gene structures of PRPS1 with PRPS2 and PRPSAP2 with PRPSAP1 revealed conserved splice site junctions, thus confirming their origins (Supplementary Figs.2a and 2b). This co-evolution pattern between PRPS and PRPSAP, indicates a potential interdependence. Additionally, PRPS1L1, a testis-restricted intronless isoform^4^, arose in the common ancestor of Eutherians presumably through a retrotransposition event involving PRPS1-encoding transcript (Fig.1c). Based on the rapid phenotypic evolution of testes^19^, we speculate that the retention of PRPS1L1 contributed to facilitate those evolutionary changes. Interestingly, in certain mammals, including mice and humans, PRPS1 and PRPS2 are located on the X chromosome, whereas PRPS1L1 is autosomal and may play a role in supporting spermatogenesis in testes.

In mammals, while PRPS1 and PRPS2 have been extensively studied in vitro for their biophysical and biochemical properties, PRPSAPs (PRPSAP1 and PRPSAP2) have not been thoroughly characterized^8–10,20–23^. Motivated by the broad conservation of PRPSAP2 across opisthokonts and its origin from PRPS1, we investigated whether PRPSAPs could independently catalyze reactions similar to PRPS isozymes. In PRPS isozymes, two subunits of a dimer form a minimal functional unit to generate the active site where the catalysis is facilitated by surrounding flexible loop regions^14,20^. The FLAG region of one PRPS monomer (Subunit A) coordinates ATP binding while the regulatory flexible loop, pyrophosphate (PP) loop, catalytic flexible (CF) loop, and ribose-5-phosphate (R5P) loop from another monomer (Subunit B) participate in catalysis (Fig.1d). A comparison of active site residues between PRPS1 and PRPSAP2 revealed a high degree of conservation in PRPS1 across Opisthokonta (Supplementary Fig.3a). In contrast, adaptive changes occurred rapidly in the loop regions of PRPSAP2 post-duplication suggesting strong selective pressure against the catalytic function. For instance, in human PRPS1, critical catalytic residues such as D171, K194, R196, N200, T225 have S177, G200, A202, E206, and D261 at corresponding positions in human PRPSAP1 (Figs.1d and 1e), preventing PRPSAPs from coordinating interactions between ATP and R5P in the closed conformation of catalytic loop, and from stabilizing the transition state^20,22,24,25^. Similarly, NHR containing homologs from Amoebozoa, Apusozoa and CRuMs also demonstrate poor conservation of active site residues in one monomer suggesting that they are unlikely to independently catalyze PRPP from R5P and ATP (Supplementary Table 1, Supplementary Fig.4a).

Notably, the two interfaces that facilitate intramolecular interactions between PRPS enzyme subunits^14,20,22,23,26^– a bent dimer which is essential for catalysis, and a parallel dimer which is essential for allostery, are highly conserved in PRPSAPs suggesting a potential regulatory mechanism involving intermolecular binding with the isozymes (Supplementary Fig.3b). The same interfaces appear to be conserved between PRPS enzymes and their non-catalytic NHR-containing paralogs in non-opisthokonts, suggestive of their ability to form heteromeric complexes (Supplementary Figs.4b-d). Collectively, the independent emergence of such homologs in both opisthokonts and non-opisthokonts within Amorphea, sharing similar patterns of conservation and divergence, strongly supports the case for convergent evolution with potential regulatory roles via interaction with corresponding PRPS isozymes in a multimeric PRPS complex.

### PRPS enzyme complex can be arranged in heterogeneous configurations

Given the conserved interaction interfaces, we experimentally tested complex formation between PRPS paralogs in mammalian cells using mouse embryonic fibroblasts (NIH3T3) and human embryonic kidney cells (HEK293T). GFP-tagging all individual paralogs followed by immunoprecipitation assays revealed interactions among PRPS1, PRPS2, PRPSAP1 (AP1), and PRPSAP2 (AP2), confirming previous studies^8–10^, and establishing the stable nature of the PRPS enzyme complex (Figs.2a and 2b, Supplementary Figs.5a-c, Supplementary Table 2 and 3). Notably, we detected the testis-specific isoform PRPS1L1 in HEK293T cells but not in NIH3T3 cells. A knock-in NIH3T3 cell line with ALFA-tagged PRPS1 confirmed these findings at endogenous expression levels as well (Supplementary Figs.5d and 5e). To further characterize this four-component complex, we employed analytical size-exclusion chromatography (SEC) to assess its molecular weight. To establish intra-run controls suitable for cross sample comparison, we developed a panel of internal standards comprised of ubiquitously expressed proteins of known native molecular weights (MWs) and validated antibodies to serve as molecular weight markers for the fractions collected (see Methods). These standards cover a wide range of MWs from 1.5 MDa to 27 kDa (smaller than monomeric PRPS1). Nearly all PRPS paralogs are involved in heteromeric associations as evident from the overlapping retention times with an estimated average complex size of ∼1.5 MDa for both NIH3T3 and HEK293T cells (Fig.2c, Supplementary Fig.5f). Given this substantial size, we wondered how the PRPS complex compared with other assemblies within a similar high molecular weight (HMW) range. A proteomic analysis of HMW protein fractions from SEC (Supplementary Fig.5g) identified a total of 262 unique proteins, which included ribosomal proteins and CAD (one of our standards), confirming enrichment for HMW proteins. This unbiased proteomic strategy revealed that among the eight cytosolic enzymes residing in HMW range, two were PRPS isozymes, indicating that the PRPS enzyme complex is one of the largest cytosolic metabolic assemblies in cells (Supplementary Figs.5h and 5i, Supplementary Table 4).

**Fig. 2:**
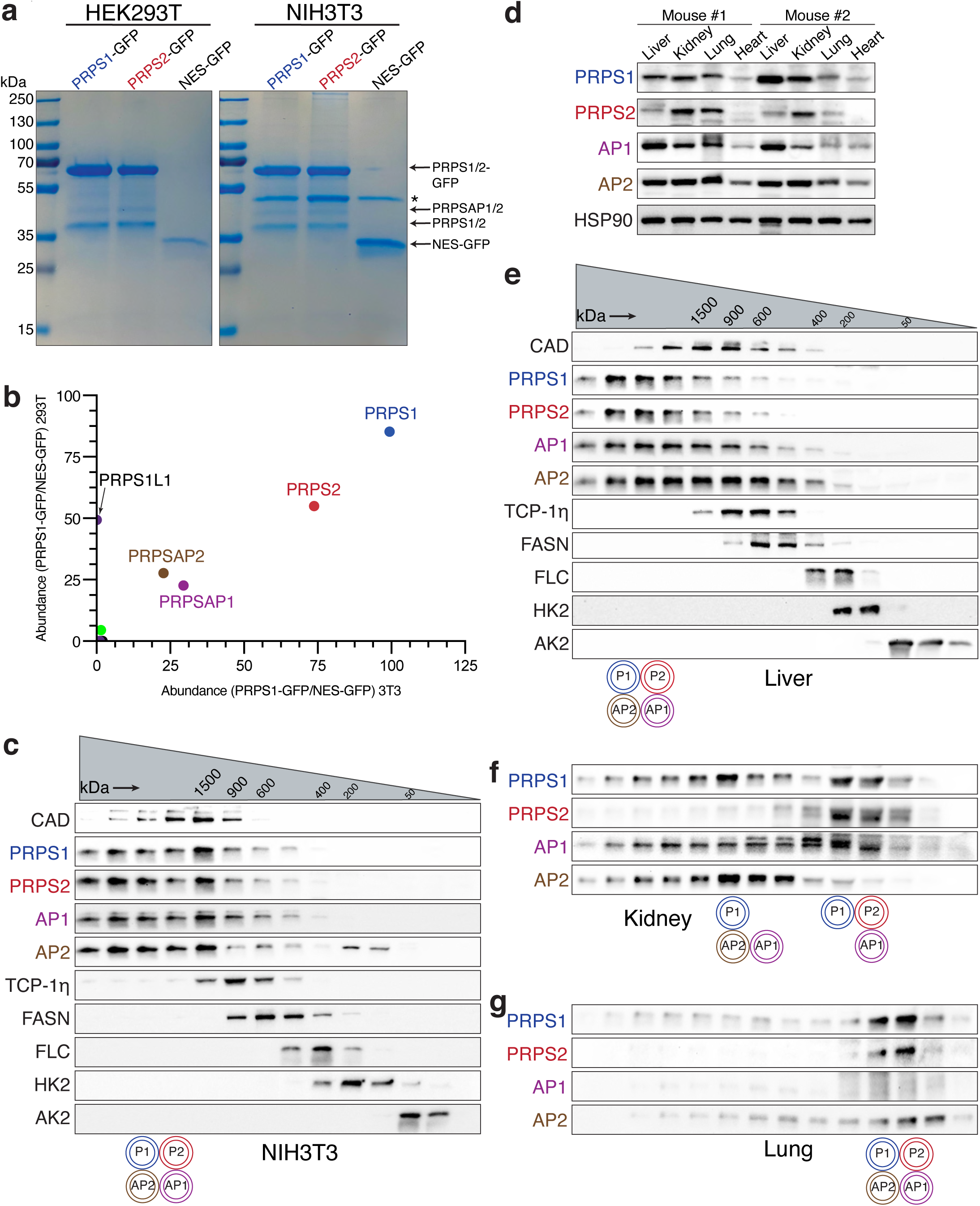
PRPS enzymes operate as a multimeric complex that attains heterogeneous configurations. a) SDS-PAGE followed by Coomassie stain of the eluates from GFP immunoprecipitation (IP) performed in NIH3T3 and HEK293T cells stably expressing PRPS1-GFP, PRPS2-GFP and NES-GFP. NES (Nuclear export signal)-GFP cells were used as controls. Asterisk indicates a non-specific band in the eluates. b) Scatter plot from mass spectrometry (MS) runs of eluates from GFP immunoprecipitation (IP) in NIH3T3 and HEK293T cells stably expressing PRPS1-GFP on the x-axis and y-axis, respectively. The axes represent the square root-transformed SEQUEST HT scores of PRPS1-GFP normalized to control. PRPS1L1 isoform was detected in HEK29T cells only. c) Western blot analysis of size exclusion chromatography (SEC) fractions collected from NIH3T3 native whole cell lysates. Cell lysates were fractionated using a Superose 6 Increase 3.2/300 column. Immunoblots probing indicated PRPS complex members and internal standards are shown. d) Protein expression profiles of PRPS complex components in various mouse tissues. HSP90 was used as a loading control. 12-week-old male C57BL/6 mice were used. e) and f) and g) represent western blot analysis of SEC fractions collected from native tissue lysates of liver, kidney, and lung, respectively. 12-week-old male C57BL/6 mice were used. Cell lysates were fractionated on a Superose 6 Increase 3.2/300 column. Immunoblots probing PRPS complex members and internal standards are shown. The circular pictograms on the bottom of the SEC immunoblots represent the schematized configurations of PRPS complex. A double circle means multiple copies of the protein are interacting within the heteromeric complex.

We next explored whether the PRPS complex exists in a similar configuration in tissues compared to our proliferating cells in culture. We observed varying expression levels for each component, indicating ubiquitous presence in different tissues (Fig.2d). Similarly, transcript levels for PRPS complex members in rat tissues (PRPS1, PRPS2, and AP1)^10,27^ and human tissues (all members) showed ubiquitous expression, albeit at different levels (Supplementary Fig.5j) perhaps indicating tissue specific catalytic and/or regulatory properties. In the liver, the PRPS complex size was ∼1.5 MDa, with most members interacting together (Fig.2e). In the kidney, multiple configurations were observed: one complex at 1 MDa with PRPS1, AP1, and AP2, and another smaller complex consisting of PRPS1, PRPS2, and AP2, which may represent cell type differences in composition (Fig.2f). In the lung, which exhibited low AP1 and AP2 expression, the complex size was the smallest among the tested tissues (Fig.2g). These results demonstrate a heterogeneous array of PRPS complex configurations, capable of forming assemblies smaller than 100 kDa and greater than a megadalton. Importantly, variations in PRPS complex architecture appear to correlate with stoichiometric expression of PRPSAPs suggestive of a crucial role in coordinating complex assembly. Functional studies have demonstrated that interactions between yeast PRPSAP-like orthologs and PRPS isozymes are essential for cell viability, further highlighting the importance of maintaining proper PRPS complex architecture^28,29^.

### Genetic knockout studies reveal severe impact on metabolism and proliferation in cells exclusively expressing PRPS1

To ascertain whether interactions between mammalian PRPS paralogs are functionally significant, we employed a CRISPR-Cas9 knockout (KO) strategy in NIH3T3 cells to generate all viable genetic knockout combinations (Fig.3a and Supplementary Fig.6a). Notably, we were unable to generate P1/AP1/AP2 KO cells, indicating that PRPS2 may not be sufficiently stable or active as a standalone enzyme in cells. Interestingly, the loss of AP1; AP2, or both associated proteins resulted in more severe cellular proliferation defects compared to the knockout of either PRPS1 or PRPS2 (Fig.3b and Supplementary Fig.6b). Of all the knockout clones tested, P2/AP1/AP2 KO and P2/AP2 KO cells showed the most pronounced proliferation defects.

**Fig. 3:**
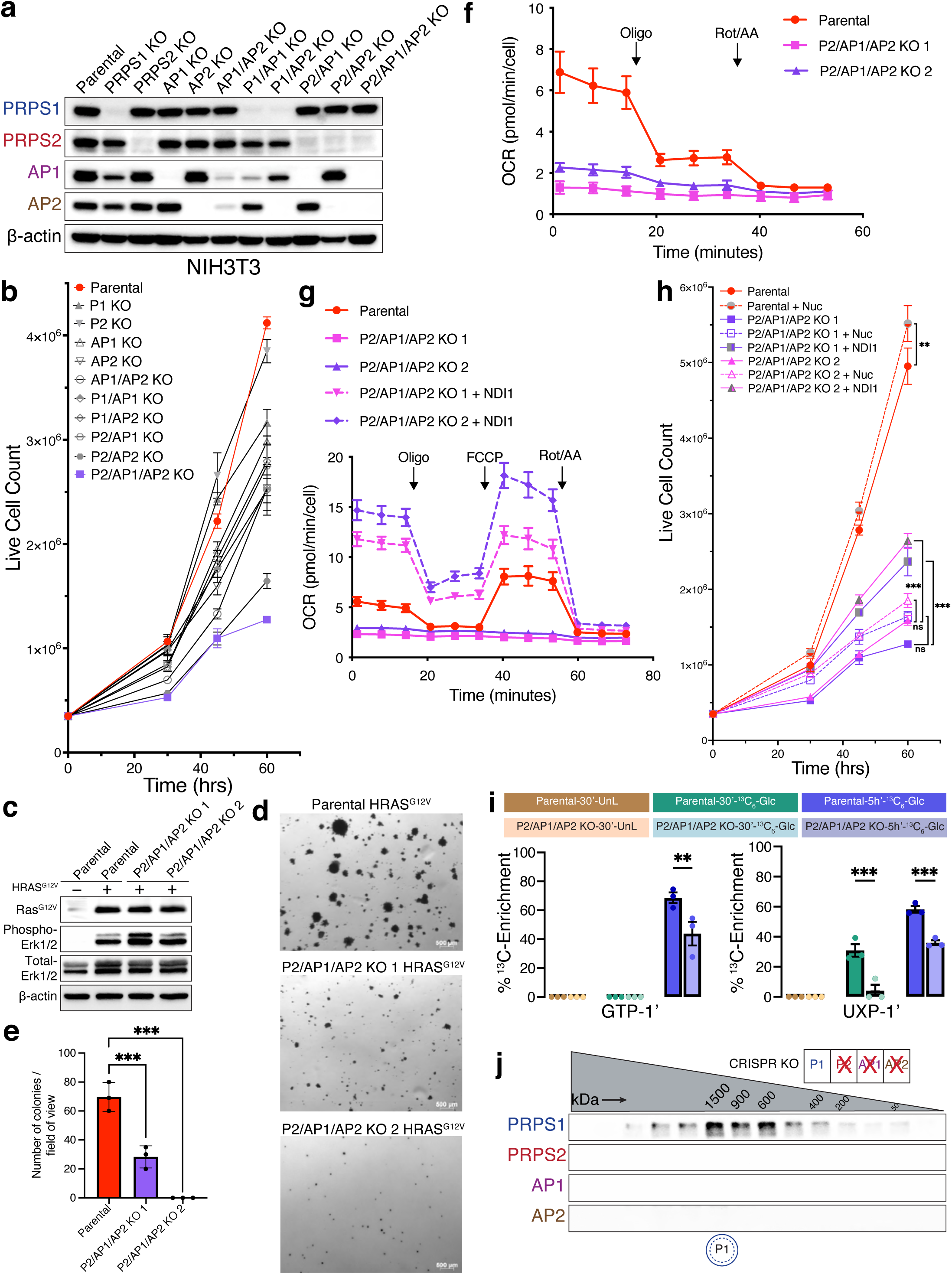
Cells exclusively expressing PRPS1 display metabolic defects and decreased cellular proliferation. a) Western blot validating CRISPR-Cas9-generated isogenic knockout cell lines. b) Proliferation for the panel of NIH3T3 knockout cell lines generated in a). c) Western blot validating HRAS^G12V^-overexpression in parental and P2/AP1/AP2 KO cell lines. Phospho-MAPK (Erk1/2) (T202/Y204) was used as a marker for activation of signaling pathways upon HRAS^G12V^ overexpression. d) Representative colony images from soft agar colony formation assay performed in NIH3T3 parental and P2/AP1/AP2 KO cells expressing HRAS^G12V^. e) Colony numbers from d). Error bars represent SD; n = 3 (***P < 0.001 by unpaired two-tailed t-test). f) Oxygen consumption rate (OCR) measured by Seahorse ATP Rate Assay in parental and P2/AP1/AP2 KO cell lines. Error bars represent SD; n = 7. g) Oxygen consumption rate (OCR) measured by Seahorse mitochondrial stress tests for NDI1-expressing P2/AP1/AP2 KO cell lines. Error bars represent SD; n = 7. h) Proliferation for nucleoside supplemented and NDI1 expressing P2/AP1/AP2 cell lines. Error bars represent SD; n = 3. (**P < 0.01; ***P < 0.001 by one-way ANOVA with post-hoc test). i) ^13^C6-glucose metabolic labeling performed in NIH3T3 parental and P2/AP1/AP2 KO cell lines for 30 minutes (30’) and 5 hours (5h’). Unlabeled, ^13^C-labeled (30 mins), and ^13^C-labeled (5 hours) sets are represented in brown, green and blue colors, respectively. ^13^C-enrichments were quantified from ^1^H-NMR spectra. Error bars represent SE; n = 3 (**P < 0.01; ***P < 0.001 by unpaired two-tailed t-test). j) Western blot analysis of SEC fractions collected from NIH3T3 P2/AP1/AP2 KO native whole-cell lysates. Cell lysates were fractionated on a Superose 6 Increase 3.2/300 column. In the pictogram, a double circle with a dotted inner circle means multiple copies of PRPS1 are forming homo-oligomers.

Collectively, these findings highlight the importance of partnerships within the PRPS enzyme complex in establishing specific configurations that are vital for maintaining optimal PRPS activity in mammalian cells.

To test whether an increased anabolic stimulus would circumvent or augment the diminished PRPS-controlled proliferative capacity, we overexpressed mutant oncogenic Ras, which is known to transform cells in part by rewiring metabolism to increase pentose phosphate pathway flux and nucleotide production^30^. Upon transduction of NIH3T3 parental cells and P2/AP1/AP2 KO cells with H-Ras^G12V^, we observed that parental cells readily formed colonies in soft agar medium, while P2/AP1/AP2 KO cells produced significantly fewer and smaller colonies suggesting PRPS activity may impose a strict ceiling on metabolic flux (Figs. 3c-e). To understand the basis for impaired cell proliferation, we first checked cell cycle profiles which revealed there was no cell cycle arrest in the knockout clones (Supplementary Fig.6c). Additionally, cleaved-PARP1 levels were comparable across parental and knockout cells, indicating that apoptosis is not induced in the knockout cells (Supplementary Fig.6d). Collectively, these results point toward a slower progression through the cell cycle in knockout cells likely due to decreased PRPP synthesis rates.

To pinpoint the metabolic basis for decreased cell cycle progression, we first measured total ATP levels in parental and P2/AP1/AP2 KO cells and observed a slight decrease in P2/AP1/AP2 KO cells (Supplementary Fig.6e). However, that modest difference was not sufficient to trigger energy stress as phosphorylation of AMPK remained consistent between parental and knockout clones (Supplementary Fig.6d). Using the Seahorse assay to evaluate mitochondrial and glycolytic energy production, we found that, despite comparable total ATP production rates, P2/AP1/AP2 KO cells had significantly lower respiration-linked ATP production rates, which were compensated by an increase in glycolytic ATP production rates compared to wild-type cells (Supplementary Fig.6f). Measuring mitochondrial oxygen consumption rates (OCR) revealed a striking loss of mitochondrial respiration in P2/AP1/AP2 KO cells, perhaps reflective of altered redox homeostasis^31^ (Fig.3f). To rescue this mitochondrial defect, we overexpressed NDI1 (yeast complex I)^32^ in P2/AP1/AP2 KO cells. NDI1 successfully increased mitochondrial OCR, including both basal and maximal rates, and enhanced ATP production (Fig.3g and Supplementary Fig.6g). Improving mitochondrial respiration offered slightly better rescue than nucleoside supplementation alone, however neither response was sufficient to restore proliferation rates to that of parental cells suggesting that P2/AP1/AP2 KO cells suffer from a more widespread metabolic defect (Fig.3h). To investigate whether a decreased flux in PRPP-consuming metabolic routes might be driving these metabolic defects including mitochondrial dysfunction in P2/AP1/AP2 KO cells, we performed ^13^C6-glucose tracing experiments, which revealed that P2/AP1/AP2 KO cells indeed have decreased pentose phosphate pathway flux and nucleotide production in addition to changes in glycolysis, creatine phosphate pathway, and choline metabolism (Fig.3i and Supplementary Table 5). These results demonstrate that PRPS1 as a standalone enzyme in cells is insufficient to sustain the metabolic flux necessary to keep up with the demands of proliferation.

Given that wild-type cells produce high molecular weight PRPS complex assemblies, we hypothesized that PRPS1 alone might be insufficient to form the large oligomers required for optimal activity. However, we observed that PRPS1 alone could still form higher-order homotypic assemblies (Fig.3j) consistent with recent cryo-EM structures showing filamentous assembly of PRPS1^22,26^. Since AP1/AP2 KO cells proliferate faster than P2/AP1/AP2 KO cells (Fig.3b and Supplementary Fig.6b), we speculated that in AP1/AP2 KO cells, PRPS1 may assemble with PRPS2 to form mixed filaments thereby enhancing the complex activity, especially since PRPS2 has been shown to form filaments in vitro as well^23^. Paradoxically, reexpressing exogenous PRPS2 in PRPS1-only (P2/AP1/AP2 KO) cells restricted PRPS1 into smaller complexes with PRPS2 indicative of an improved activity over PRPS1 homooligomers (Supplementary Fig.6h). Altogether, our findings suggest that homotypic PRPS1 assemblies, which can be disrupted by PRPS2, are aberrant and suboptimal configurations for cells.

### Organizing principles of PRPS complex assembly

Based on our findings that homo-oligomerization of PRPS1 is detrimental to cellular metabolism, we hypothesized that P2/AP2 KO cells, which share similar proliferative defects with P2/AP1/AP2 KO cells (Fig.3b and Supplementary Fig.6b), might contain such aberrant homotypic assemblies. Indeed, in P2/AP2 KO cells, PRPS1 and AP1 did not bind to each other; instead, they formed homo-oligomers of distinct sizes (Fig.4a), a result that was recapitulated by overexpressing AP1 in PRPS1-only cells (Supplementary Fig.7a). We further confirmed these results by performing PRPS1 pulldown in P2/AP2 KO cells where we observed minimal interactions between PRPS1 and AP1 (Supplementary Fig.7b). Surprisingly, these results demonstrate that, despite the strict conservation of dimer interfaces between PRPS isozymes and associated proteins, there may be a preferential selection of binding partners within the complex.

**Fig. 4:**
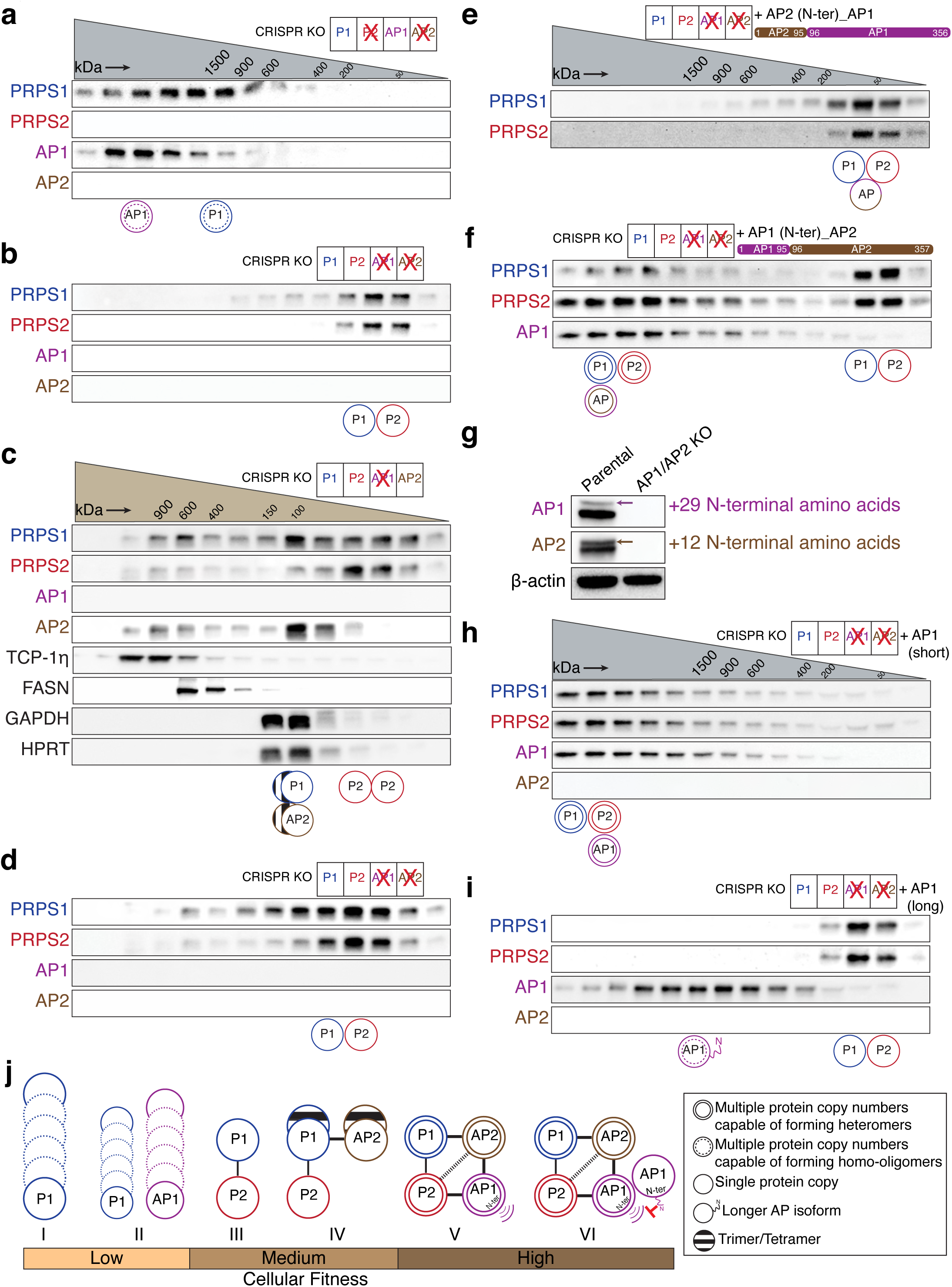
PRPSAPs are molecular scaffolds with preferential binding partners among PRPS isozymes. a-f) Western blot analysis of SEC fractions collected from native whole-cell lysates of NIH3T3 P2/AP2 KO cells (a), AP1/AP2 KO cells (b), AP1 KO cells (c) AP1/AP2 KO cells (d), AP1/AP2 KO cells stably expressing chimeric AP1 with AP2’s N-terminus (residues 1-95) (e) and chimeric AP2 with AP1’s N-terminus (residues 1-95) (f). g) Multiple isoforms for AP1 and AP2 were detected under optimal SDS-PAGE resolution. The arrows in immunoblots indicate the longer isoforms of AP1 and AP2, which are N-terminal leader sequences of 29 and 12 amino acids, respectively. h) and i) Western blot analysis of SEC fractions collected from NIH3T3 AP1/AP2 KO cells stably expressing the short isoform of AP1 (h) and long isoform of AP1 (i). Cell lysates for (a), (b), (e), (f), (h), and (i) were fractionated on a Superose 6 Increase 3.2/300 column whereas cell lysates for (c) and (d) were fractionated on a Yarra SEC-2000 column. The circular pictograms at the bottom of the SEC immunoblots illustrate the different configurations of the PRPS complex. A double circle means multiple copies of the protein are interacting within the heteromeric complex. A double circle with a dotted inner circle means multiple copies of the protein are forming homo-oligomers. A single circle means a single protein is interacting within the complex. A circle with lines inside indicates that the proteins may be forming a trimer or tetramer. j) Association of different PRPS complex configurations with cellular fitness. (I) PRPS1 alone undergoes self-assembly to form homo-oligomers. (II) PRPS1 and AP1 cannot interact directly and form separate homo-oligomers. (III) PRPS2 binds PRPS1 to form a dimer and disrupts PRPS1 homotypic assemblies. (IV) AP2 preferentially binds with PRPS1 to form a trimer/tetramer to nucleate the complex. (V) AP1 preferentially binds with AP2 and PRPS2 to elongate the PRPS complex via its N-terminus. (VI) The longer isoform of AP1 with flexible N-terminal extension likely caps the complex. The weight of the connectors between the circles representing proteins reflects the predicted strength of protein-protein interactions between them. The heavier the line, the stronger the predicted interaction between the members of the complex.

Building on the concept of specific partnerships, we next investigated the preferential binding modalities within the PRPS complex. In P2/AP1 KO cells, PRPS1 and AP2 together formed higher-order structures (Supplementary Fig.7c) resembling kidney – which exhibited low AP1 expression – where PRPS1 appeared to preferentially bind with AP2 (Fig.2f). Next, we asked whether PRPS2 and AP1 can similarly interact with each other. Indeed, in P1/AP2 KO cells, PRPS2 and AP1 formed higher-order heterotypic assemblies (Supplementary Fig.7d). This suggests a defined preferential binding between PRPS1 and AP2 – partners that have co-evolved since the early opisthokonts – as well as between PRPS2 and AP1, which originated together in the common ancestor of jawed vertebrates.

To identify the primary determinant of formation of competent HMW PRPS complex, we characterized complex assembly potential in remaining isogenic knockout series. We found that the loss of PRPS1, PRPS2 or AP2 alone did not significantly alter the formation of higher order heterotypic assemblies (Supplementary Figs.7e-g). Interestingly, in the absence of AP1 – observed in both AP1 KO and AP1/AP2 KO cells – we detected a significantly smaller PRPS complex, indicating a major role for AP1 in governing HMW complex formation (Fig.4b, Supplementary Fig.7h). In line with this, we noted that in P1/AP1 KO cells, PRPS2 and AP2 were unable to assemble into a HMW complex (Supplementary Fig.7i). Altogether, we attribute stimulation of PRPS complex elongation as an emergent property of AP1.

To investigate whether AP1 KO and AP1/AP2 KO cells form distinct PRPS complex configurations, we employed Yarra SEC-2000 column (fractionation range – 1 kDa to 300 kDa, better for smaller MW) as Superose 6 Increase 3.2/300 column (fractionation range – 5 kDa to 5 MDa) produced identical elution profiles for both knockout lines. In AP1 KO cells, PRPS1 preferentially binds with AP2 to form heterotrimers or heterotetramers, while PRPS2 primarily forms homodimers (Fig.4c). In AP1/AP2 KO cells, we predominantly observe PRPS1-PRPS2 heterodimers (Fig.4d) reminiscent of lung, which exhibited low AP1 and AP2 expression. This finding aligns with our previous results showing that reintroducing PRPS2 to PRPS1-only cells restricts PRPS1 homo-oligomerization (compare Fig.4d with Supplementary Fig.6h). Here, we demonstrate that AP2 outcompetes PRPS2 for binding to PRPS1, likely because the PRPS1-PRPS2 complex must form a bent dimer for catalysis, preventing the assembly of a parallel dimer required for allosteric site formation, which renders it insensitive to feedback regulation. In contrast, the PRPS1-AP2 complex, capable of forming either a heterotrimer or heterotetramer, can assemble both parallel and bent dimers, thereby remaining responsive to allosteric regulation.

Because disordered NHRs are hallmark features of PRPSAPs, and intrinsically disordered regions (IDRs) are known to enable biomolecular condensate formation^33^, we hypothesized that the NHR unique to PRPSAP1 might promote PRPS complex elongation. To test this, we overexpressed an NHR-deletion construct of AP1 in AP1 KO cells, which fully restored the formation of higher order heterotypic PRPS complex (Supplementary Fig.8a). These results indicate that NHRs, a feature shared between AP1 and AP2 isoforms, do not play a direct role in facilitating complex assembly. However, the positive selection of NHRs in PRPSAPs suggests that they may influence the flexibility or regulation of the complex in other ways.

To identify the structural elements that distinguish AP1 from AP2 and confer AP1 its complex elongation properties, we generated four chimeric constructs by swapping regions between AP1 and AP2 and overexpressed them individually in AP1/AP2 KO cells (Supplementary Fig.8b). Only the chimera with N-terminus of AP2 (residues 1–95) failed to restore the formation of higher order heterotypic PRPS complex, whereas the other three constructs successfully formed HMW PRPS complex, confirming the N-terminus of AP1 as the distinguishing feature that promotes HMW complex assembly (Fig.4e and Supplementary Figs. 8c-e). As expected, replacing AP2’s N-terminus with the corresponding residues from AP1 promoted the formation of larger multimers (Fig.4f). Notably, excluding the flexible residues within the NHRs, the N-terminus (residues 1–95) accounts for over 50% of the amino acid differences between human AP1 and AP2 (Supplementary Fig.8f). A comparative analysis of the N-termini across jawed vertebrates revealed that these differences have been conserved since the divergence of AP1 from AP2 (Supplementary Fig.8g). This highlights how rapid adaptive changes at the N-terminus of AP1 facilitate complex elongation, possibly reflecting an evolutionary response to the co-emergence of PRPS2 that restricts PRPS1 elongation.

While analyzing the N-terminal residues of AP1 and AP2, we noticed that the transcripts of AP1 and AP2 include an upstream alternative translation start site (TSS), which encodes an additional 29 and 12 amino acids at the N-terminus for murine AP1 and AP2, respectively (Fig.4g and Supplementary Fig.9a). These upstream sequences with potential alternative start sites for AP1 and AP2 have been positively selected for since their emergence in Osteichthyes and Amniota, respectively (Supplementary Figs.9b-d). To understand the influence of long and short isoforms, we introduced them individually into AP1/AP2 KO cells and performed SEC. Interestingly, exogenous expression of the short AP1 isoform completely restored the complex size whereas the longer AP1 isoform failed to interact with and incorporate PRPS1 and PRPS2 into higher-order structures (Figs.4h and 4i). Notably, the long AP1 isoform could undergo homo-oligomerization. In contrast, both long and short isoforms of AP2 failed to restore the complex size (Supplementary Figs.10a and 10b). The short AP2 isoform was bound with both PRPS1 and PRPS2 while the long AP2 isoform seemed to have higher affinity for PRPS1 compared to PRPS2. These results conclusively identify the short AP1 isoform as the primary driver of PRPS complex elongation and highlight the regulated assembly of PRPS complex via translational control, mediated by the inclusion or exclusion of N-terminal leader sequences.

## Discussion

Eukaryogenesis involved expansion in cell volume, genome size, number of protein-coding genes and regulation of gene expression, along with the addition of metabolically demanding compartmentalized machineries, all fueled by an energy boost from mitochondria^34^. The complexity of eukaryotes included the diversification of metabolic enzymes, likely providing the Last Eukaryotic Common Ancestor (LECA) with an adaptive advantage to inhabit various ecological niches and increasing the likelihood of survival for diverse lineages. Here, we showed that acquisition and maintenance of multiple PRPS homologs is a key trait of eukaryotic metabolic system, underscored by our demonstration that PRPS1 homo-oligomers do not function effectively to sustain eukaryotic metabolism. It is clear from our eukaryote-wide survey of PRPS homologs that later symbiotic events, genetic reshuffling, and gene duplications have provided fertile ground for sculpting a PRPS armament enriched with evolutionary innovations to enable species to adapt their metabolism. Our case study on the origins of the mammalian PRPS complex demonstrate how a single PRPS gene, over vast evolutionary timescale, ultimately gives rise to multiple discrete homologs while maintaining intermolecular interactions to create a tunable multi-component metabolic assembly (Fig.5a)^35^.

**Fig. 5:**
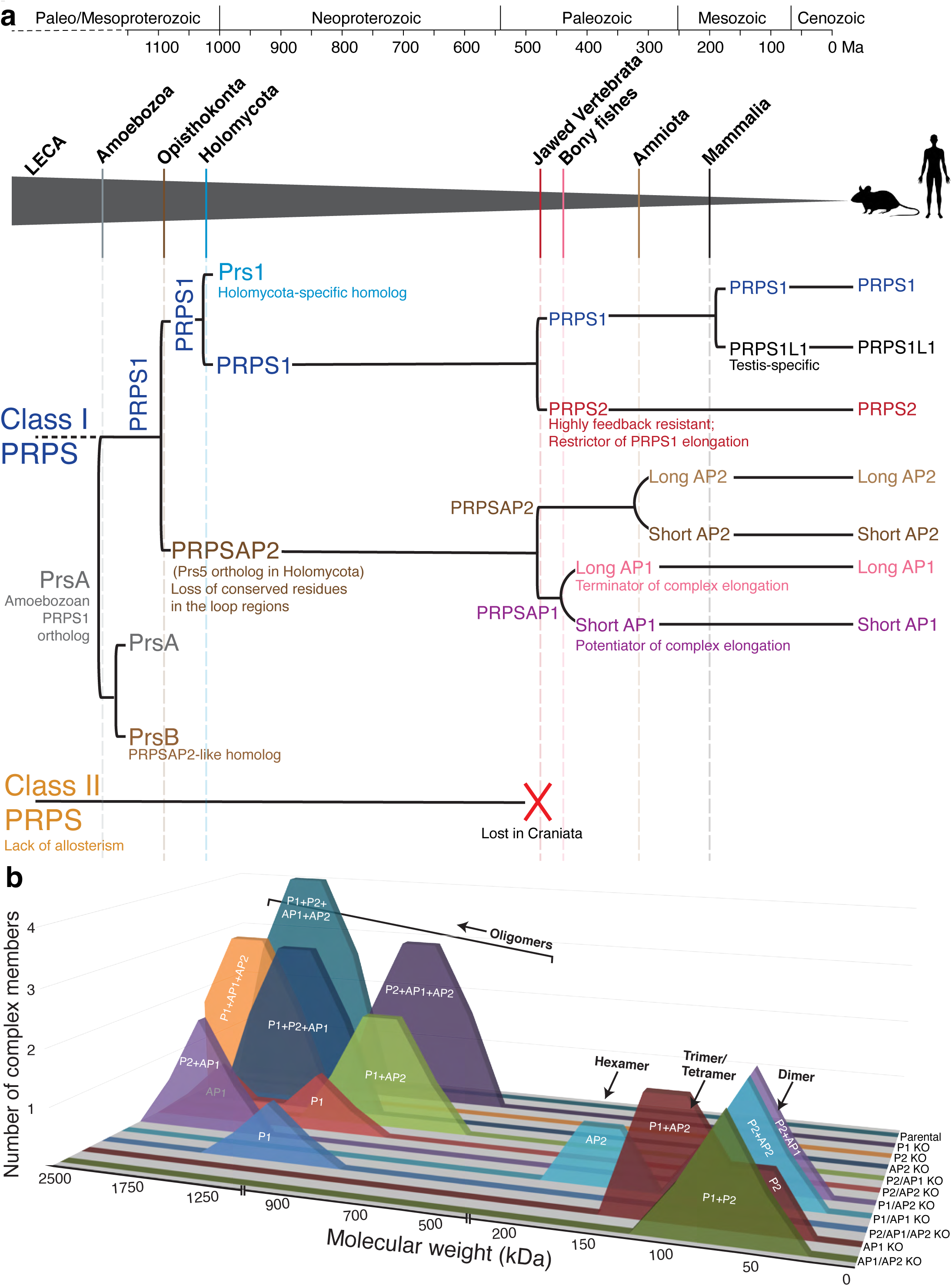
Timeline of PRPS complex evolution in mammals and summary of PRPS complex across isogenic knockout. a) Chronological duplication events of a Class I PRPS encoding gene in indicated eukaryotes. The grey triangle traces the evolutionary trajectory leading to the emergence of mammalian PRPS homologs. Solid vertical lines within the triangle denote the origin of major lineages, while dashed lines indicate significant duplication events that gave rise to PRPS homologs, with notable functional innovations occurring post duplication. Class II PRPS which was likely present in the LECA is lost in the craniates, followed by rapid expansion of Class I PRPS homologs in jawed vertebrates. Left square bracket indicates a gene duplication event and left round bracket indicates an extension at the N-terminus. The molecular clock-based evolutionary timeline (in millions of years before present, Ma) is adapted from Dohrmann et. al^35^. b) Summary of SEC profiles from NIH3T3 parental and panel of CRISPR KO lines. The x-axis represents the molecular weight, and the y-axis represents the number of PRPS complex members interacting in the complex.

The precise positive and negative selection pressures that shaped PRPS homologs after their emergence are hard to determine post hoc, but some functional changes may offer insights. Despite losing key active site residues necessary for independent PRPP generation, the conserved FLAG region (Supplementary Fig.3a) in PRPSAPs and PRPSAP-like orthologs suggests they might still form a minimal functional unit—a dimer—with a PRPS isozyme.

Interestingly, we have shown that independent gene duplications throughout evolution have enabled remodeling of flexible loops, as seen in the expansion of regulatory loops (in Prs5 orthologs) and CF loops (in PRPSAP2/PrsB/PRPSAP-like orthologs), and positive selection has preserved such innovations. An analysis of more than 30,000 proteins has shown that nature relies on a limited repertoire of basic structures (domains, motifs, and folds) to perform a wide range of functions^36^. Enzymes achieve functional innovations by altering the flexible loop structures exposed on their surfaces^37,38^, including those in PRPP-utilizing enzymes^39^. In the PRPS complex, these disordered CF loops could be primed by allosteric mechanisms and post-translational modifications (PTMs), allowing them to act as dynamic switches that can rapidly adapt to the cell’s metabolic needs by sampling various conformational states. For instance, the movement of the CF loop in PRPSAP1/2 could influence the kinetics of catalytic loop opening in adjacent PRPS1/2^22^. Different loop conformational substates might influence access to allosteric and PTM sites in the neighboring PRPS isozymes^40–42^, as well as modulate intra and intermolecular interactions mediated via allostery^2^. Additionally, PRPSAPs exhibit poor conservation in the regulatory flexible loop residues that are essential for formation of allosteric sites I and II (Subunit B in Supplementary Figs.11a-d)^14,20^. The extent of PRPSAP-mediated regulation likely depends on the specific subunit arrangement in a PRPS-PRPSAP dimer/trimer and even across protomers of a filament, as filament interfaces are highly conserved in PRPSAPs, similar to PRPS isozymes^22,23^ (Supplementary Figs.12a and 12b). Our findings pave the way for future structural and biochemical studies aimed at deciphering the molecular underpinnings of different heterogenous PRPS complex configurations.

Our phylogenomic analysis uncovered a coevolutionary pattern between Class I PRPS homologs (“classical” PRPS) and Class II PRPS enzymes, which exhibit poorly conserved dimer interfaces – both bent and parallel – and lack the allosteric regulation observed in Class I PRPS^2^. Ancestral opisthokonts and amoebozoans possessed Class II enzymes, while holomycotans lost them, concomitantly expanding their class I PRPS repertoire (Supplementary Fig.13).

Conversely, most holozoans retained Class II enzymes but possess only two Class I homologs. Similarly, loss of Class II enzymes in Cyclostomata was followed by emergence of PRPS2 and AP1 isoforms in jawed Vertebrata. These evolutionary trends may represent compensatory selection, explaining the observed expansion of Class I PRPS homologs to combat loss of allosterically resistant Class II PRPS (Supplementary Figs.11a-d), while also co-opting for evolutionary innovations. Indeed, unlike PRPS1, PRPS2 is highly feedback-resistant^43^, and we have shown that it also suppresses formation of supramolecular assemblies with PRPS1, adding another dimension to PRPS2’s ability to stimulate PRPP production. We propose that AP1 concurrently emerged to counteract PRPS2 by preferentially binding to it and restoring formation of higher-order structures, reminiscent of PRPS1’s abilities. Additionally, the duplication of AP1 from AP2 led to significant structural changes, particularly at the N-terminus, enabling it to drive higher-order heterotypic assemblies. The N-terminus is crucial, as evolution has introduced N-terminal leader sequences in both AP1 and AP2, further diversifying their roles. Similar to Myc-driven translational regulation of PRPS2^44^, translation control emerges as a crucial driver of PRPS complex assembly via TSS selection in PRPSAPs providing an important link between major anabolic sinks and metabolic flux determining enzymes, however the mechanisms determining TSS selection and physiological settings under which this regulation is important remains unknown.

Mammalian metabolic complexes are known to exhibit various molecular forms to tailor enzymatic profiles according to metabolic demands^45^. Our discovery that the PRPS complex alters its architecture across different tissues, combined with recent evidence of varied purine synthesis rates across tissues^46^ where PRPS is a major determinant of purine production^47–49^, suggests that the flexible nature of the PRPS complex is vital for metabolic adaptation. The specific impact that PRPS complex modulation has on cellular biochemistry may be cell-type specific^50^. Our genetic knockout studies in cells support the existence of multiple configurations of PRPS complex, which can result in varying levels of cellular fitness (Figs.4j and 5b). These myriad assembly states, involving changes in stoichiometry, topology, dynamic conformations, and differential affinities, have significant implications for regulatory and catalytic properties.

Our research lays the groundwork for future studies focused on understanding the mechanisms underlying the upstream signals that regulate PRPS complex assembly and the associated activity and metabolic output of specific PRPS complex configurations. Ultimately, this study advocates for a paradigm shift, highlighting that PRPS activity is regulated at the level of the complex— shaped by over a billion years of evolution— which may explain the variable penetrance and pleiotropic effects of idiopathic PRPS disease mutations.

## Methods

### Database search and annotation for sequences

Amino acid sequences from well annotated PRPS enzymes from model organisms were used as templates to identify homologs in the NCBI database using BLAST^51^. Hits were subsequently curated manually based on significant matches, protein domains, and conserved motifs.

For manual annotation, datasets were retrieved from sequence read archive (SRA)^52^. Selection criteria prioritized datasets with high-quality sequencing reads and comprehensive metadata, ensuring suitability for downstream protein annotation. Quality control checks were performed using BlastP versus eukaryotic or prokaryotic domains. Open reading frames (ORFs) were identified within transcripts, and potential protein sequences were deduced based on canonical start and stop codons, as well as sequence homology to well-annotated PRPS proteins.

For splicing analysis, exonic and intronic sequences were sourced either from NCBI/Ensembl^53^ databases or manually curated from indicated genomic sequences, taking into account splice junctions rules (canonical GT/AG and non-canonical GC/AG). Where available splice prediction was corroborated with transcriptomics of the same species or nearest available relative.

Multiple sequence alignment (MSA) was conducted using Clustal Omega^54^ or MUSCLE^55^. A sequence logo representing desired regions was generated using the WebLogo^56^ tool online, based on the MSA results. Multiple logos were aligned using MetaLogo^57^.

### PRPS1 structure generation

The PRPS1 crystal structure with the ID – 2hcr was extracted from the PDB. Missing residues in the N-terminus, C-terminus and catalytic flexible regions were modeled using MODELLER^58^ in UCSF ChimeraX^59^. Multiple loop models were generated, and the final selections were based on the lowest normalized discrete optimized protein energy score (zDOPE), where a negative score denotes better predictions.

### AlphaFold2 structure predictions

AlphaFold modelling^60^ was performed using the Alphafold2_mmseq2^61^ notebook which was run on Google Collaboratory cloud^62^. AlphaFold’s top-scoring models were ranked from 1 to 5 by per-residue Local Distance Difference Test (pLDDT) scores (a per-residue estimate of the prediction confidence on a scale from 0 – 100). The model with the highest average pLDDT scores was used to assess the predicted structures. The selected model was visualized and analyzed using ChimeraX.

### Cell lines and cell culture conditions

NIH-3T3 cells were cultured in Dulbecco’s modified Eagle medium (DMEM) containing 10% (v/v) fetal bovine serum (FBS) and 1x penicillin/streptomycin.

### Cell growth assay

Cells were seeded onto a 6 cm plate at a density of 350,000 cells/plate. Cells counts were counted at three time points: 30 hours, 45 hours, and 60 hours after seeding. Cell counting was performed using trypan blue staining and a hemocytometer.

### Plasmids and transfection

*Prps1* cDNA (Horizon Discovery #MMM1013-202859297), *Prps2* cDNA (previously described^44^), and *PRPSAP1* cDNA (Horizon Discovery #MHS6278-202757585) were subcloned into the pMSCV-puro vector, which harbors GFP at the C-terminus, using the SalI and EcoRI sites. *PRPSAP2* cDNA (Horizon Discovery #MHS6278-202759946) was subcloned into the pMSCV vector with GFP at the C-terminus using Gibson assembly.

*PRPSAP1* and *PRPSAP2* cDNA from Horizon Discovery encode the short PRPSAP1 isoform and the long PRPSAP2 isoform, respectively. Construct encoding yeast *NDI1* retroviruses (Addgene #72876) has been previously described^63^. *Myc* cDNA in pWZL Blast myc (Addgene #10674) was replaced with *H-Ras V12* cDNA from pBABE-puro H-Ras V12 (Addgene #9051) to generate pWZL H-Ras V12 blast.

Lentiviral expression plasmids for PRPS complex members were derived as follows. First, the blasticidin expression cassette from pWZL Blast myc (Addgene #10674) was subcloned into FUGW_bleo (Addgene # 14883) to replace bleomycin. All four cDNAs encoding PRPS1, PRPS2, PRPSAP1, and PRPSAP2 were PCR amplified from pMSCV-puro vector and ligated with PCR-amplified FUGW_blasticidin backbone using Gibson assembly. Sequences encoding the ALFA epitope tag were engineered in the primers to C-terminally tag the proteins. To generate the long PRPSAP1 isoform, sequences encoding the additional 29 amino acids were ordered as gBlocks gene fragments from IDT and subcloned into FUGW_PRPSAP1-ALFA vector using Gibson assembly. The short isoform of PRPSAP2 was subcloned from FUGW_PRPSAP2-ALFA vector using Gibson assembly. To create the PRPSAP1(1-95)_PRPSAP2-ALFA construct, the first 95 amino acids of PRPSAP1 were PCR amplified from the FUGW_PRPSAP1-ALFA template. Using Gibson assembly, this amplified fragment was then ligated with the PCR-amplified FUGW_PRPSAP2-ALFA construct, which had its first 95 amino acids from PRPSAP2 removed. Other chimeric mutants were generated using similar Gibson assembly approach. Site Directed Mutagenesis (SDM) was performed to generated PRPSAP1 construct lacking non-homologous region (NHR).

Lipofectamine 3000 Transfection Reagent (Invitrogen #L3000001) was used to transfect NIH3T3 cells according to manufacturer’s instructions.

### Virus production and transduction

Retrovirus was produced in HEK293T cells by co-transfecting the retroviral transfer vector with pUMVC (Addgene #8449) and pMD2.G (Addgene #12259). For lentivirus production, the lentiviral transfer vector was used in conjunction with psPAX2 (Addgene #12260) and pMD2.G (Addgene #12259). Transfection was carried out using PolyFect transfection reagent (QIAGEN #301105), and the media was changed after 24 hours.

The supernatant containing the virus was collected at 48 hours and 72 hours post-transfection, filtered, and concentrated using Retro-X (Takara #631456) or Lenti-X (Takara #631232) concentrators. The resulting viral pellet was resuspended in fresh cell growth media and added to target cells. After 24 hours of infection, the virus was removed, and the cells were provided with appropriate selection media.

### Mice

Male C57BL/6 mice, 12 weeks of age, were used to extract tissues for size exclusion chromatography (SEC) experiments.

### Tissue protein extraction

After anesthesia, the mice were transcardially perfused with PBS through the ventricular catheter. Organs/tissues were harvested and snap frozen in liquid nitrogen, then homogenized using a mortar and pestle. For western blotting, samples were lysed with RIPA buffer (Thermo #89901) containing 1X protease and phosphatase inhibitor cocktail (Thermo #78446) while sample for SEC experiments were lysed in non-denaturing lysis buffer (50 mM Tris-Cl, pH 7.5, 200 mM NaCl, 1% digitonin, 1mM TCEP, 1mM MgCl2, benzonase and 1X protease and phosphatase inhibitor cocktail).

### SDS-PAGE and Western Blotting

For denaturing cell lysis, cells were first rinsed once with ice-cold PBS and then lysed in RIPA buffer (Thermo Scientific #89901) supplemented with 1X protease and phosphatase inhibitor cocktail (Thermo #78446). The cleared protein lysates were subsequently mixed with 1X Laemmli sample and separated on either 10% or 12% TGX Fastcast gels (BioRad #1610173 and #1610175). The proteins were then blotted onto 0.2 µm PVDF membrane using Bio-Rad Trans-Blot Turbo Transfer system. PVDF membranes were blocked for 40 mins at room temperature (RT) with 5% (w/v) milk in Tris-Buffered Saline (TBS) with 0.1% Tween-20 (TBS-T). After blocking, the membranes were washed and incubated overnight at 4°C with primary antibodies (diluted 1:1000) prepared in 3% BSA in TBS-T. Subsequently, the membranes were washed again and incubated with corresponding secondary antibodies from Jackson ImmunoResearch (diluted 1:25000) in 5% (w/v) milk in TBS-T. Blots were visualized using chemiluminescent substrates from Thermo on ChemiDoc Imaging System (BioRad #12003153). To facilitate re-probing, Restore PLUS Western Blot Stripping Buffer (Thermo #46430) was used to strip the blots.

### Antibodies

Primary antibodies used were: CAD (Cell Signaling #93925), TCP1-η (Santa Cruz #sc-271951), FASN (Cell Signaling #3180), FLC (Santa Cruz #sc-390558), HK2 (Cell Signaling #2867), AK2 (Santa Cruz #sc-374095), PRPS1/2 (Santa Cruz #sc-100822), PRPS1 (Proteintech #15549-1-AP), PRPS2 (Sigma #SAB2107995), PRPS1/2/3 (Santa Cruz #sc-376440), PRPSAP1 (Santa Cruz #sc-398422), PRPSAP2 (Proteintech #17814-1-AP), HSP90 (Cell Signaling #4877), β-Actin (Cell Signaling #4970; Cell Signaling #3700), ALFA-HRP (SynapticSystems # N1505-HRP), XO (Abcam #109235), Ras (G12V Mutant Specific) (Cell Signaling #14412), Phospho-p44/42 MAPK (Erk1/2) (Thr202/Tyr204) (Cell Signaling #4376), p44/42 MAPK (Erk1/2) (Cell Signaling #9102), β-Tubulin (Cell Signaling #2128), Phospho-AMPKα (Thr172) (Cell Signaling #2535), AMPKα (Cell Signaling #2532), cleaved PARP1 (Abcam #32064), GAPDH (Cell Signaling #5174), HPRT (Abcam #109021).

Secondary Antibodies used were: Anti mouse (Jackson ImmunoResearch #115-035-003), Anti rabbit (Jackson ImmunoResearch #111-035-003).

### GFP Immunoprecipitation Assay

Cells stably expressing GFP tagged proteins were harvested and cell pellets were lysed in non-denaturing lysis buffer (50 mM Tris-Cl, pH 7.5, 200 mM NaCl, 1% digitonin, 1mM TCEP, 1mM MgCl2, benzonase and 1X protease and phosphatase inhibitor cocktail) for 20 mins on ice. The cell lysates were then clarified by centrifugation (15,000 × g, 15 mins, 4 °C) and incubated with the equilibrated anti-GFP nanobody conjugated to magnetic particles (Chromotek GFP-Trap® Magnetic Particles M-270) for 1 h on a rotator disk at 4 °C. The supernatant was removed, and the beads were washed four times in wash buffer (10 mM Tris-Cl, pH 7.5, 1.5 M NaCl, 0.5% Triton X-100, 0.5 mM EDTA). The beads were then boiled for 10 mins at 95°C in 2X SDS-sample buffer for protein elution. The eluates were sent to University of Cincinnati Proteomics Laboratory (UCPL) for mass spectrometry (MS) analysis.

### ALFA-tag Pulldown Assay

NIH3T3 cells endogenously expressing PRPS1-ALFA^64^ proteins were lysed using non-denaturing lysis buffer (50 mM Tris-Cl, pH 7.5, 200 mM NaCl, 1% digitonin, 1mM TCEP, 1mM MgCl2, benzonase and 1X protease and phosphatase inhibitor cocktail) for 20 mins on ice. The cell lysates were then clarified by centrifugation (15,000 × g, 15 mins, 4 °C) and incubated with the equilibrated anti-ALFA nanobody conjugated to agarose beads (ALFA SELECTOR ST, Nanotag #N1511) for 1 h on a rotator disk at 4 °C. The supernatant was removed, and the beads were washed four times in wash buffer (25 mM Tris-Cl, pH 7.5, 0.5 M NaCl, 0.5% Triton X-100). Finally, the beads were boiled for 10 mins at 95°C in 2X SDS-sample buffer for protein elution.

### Size Exclusion Chromatography

Cells or tissues were lysed using non-denaturing lysis buffer (50 mM Tris-Cl, pH 7.5, 200 mM NaCl, 1% digitonin, 1mM TCEP, 1mM MgCl2, benzonase and 1X protease and phosphatase inhibitor cocktail) for 20 mins on ice. The cell lysates were then clarified by centrifugation at 15,000 × g for 15 mins at 4 °C and subsequently filtered using a 0.22 µm. About 200 μg of cell lysates were loaded onto either a Superose 6 Increase 3.2/300 column (GE Healthcare #29-0915-98) at the flow rate of 0.04 mL/min or Yarra 3μm SEC-2000 at the flow rate of 0.5 mL/min (Phenomenex #00H-4512-K0) using Thermo Vanquish UHPLC.

After passing through the void volume, the sample fractions were collected and concentrated using a 3K MWCO filter (Thermo #88512) and analyzed via western blotting. Gel filtration calibration kits (GE #28-4038-41, GE #28-4038-42, Sigma #MWGF200-1KT) were used to monitor column performance over time. Different internal standards were probed for molecular weight calibration: CAD (GLN-dependent carbamoyl phosphate synthetase (<u>C</u>PS-2), aspartate transcarbamoylase (<u>A</u>TC), and dihydroorotase (<u>D</u>HO))^65^, TCP-1η (T-Complex Protein 1 subunit eta)^66^, FASN (Fatty Acid Synthase)^67^, FLC (Ferritin Light Chain)^68^, XO (Xanthine Oxidase)^69^, GAPDH (Glyceraldehyde 3-phosphate dehydrogenase)^70^, and HPRT (Hypoxanthine phosphoribosyltransferase)^71^ form complexes of around 1500 kDa, 900 kDa, 540 kDa, 480 kDa, 290 kDa, 144 kDa, 100 kDa respectively, while HK2 (Hexokinase 2) and AK2 (Adenylate Kinase 2) are mostly monomeric at 102 kDa and 27 kDa, respectively. The elution profile of PRPS was compared with that of known globular protein complexes, which were used as internal standards to estimate the size of the PRPS enzyme complex. Internal standards were probed in every SEC run.

### Mass Spectrometry Analyses

To analyze fractions collected from SEC of HEK293T cells, cells were first lysed using non-denaturing lysis buffer (50 mM Tris-Cl, pH 7.5, 200 mM NaCl, 1% digitonin, 1mM TCEP, 1mM MgCl2, benzonase and 1X protease and phosphatase inhibitor cocktail) for 20 mins on ice. The cell lysates were then clarified by centrifugation at 15,000 × g for 15 mins at 4 °C and subsequently filtered using a 0.22 µm. About 200 μg of cell lysates were loaded onto Bio SEC-5 2000Å (Agilent #5190-2543) at the flow rate of 0.2 mL/min using Thermo Vanquish UHPLC. Samples were pooled from the indicated fractions and concentrated with a 10K MWCO filter (Thermo #88517). Subsequently, the samples were dried in a speed vacuum centrifuge and resuspended in TEAB buffer according to a standard in-solution digestion protocol. The samples were reduced with TCEP (tris-(2-carboxyethyl) phosphine) followed by alkylation with MMTS (methyl methanethiosulfonate). The samples were digested overnight at 37°C and was stopped by adding 10% formic acid. After drying, the samples were resuspended in 20 μl of 0.1% formic acid. A quarter of each sample was then subjected to nanoLC-MS/MS analysis (Orbitrap Eclipse). Peptide fragmentation spectra were searched against the human database using Proteome Discoverer version 2.4 and the Sequest HT search algorithm (Thermo).

For the analysis of eluates from GFP immunoprecipitation, 30 μL of each eluate sample was loaded onto Invitrogen 4-12% Bis-Tris gels and separated using MOPS buffer. Pre-stained MW marker were used between lanes to facilitate cutting out the full protein region for each sample. The bands were excised, reduced with DTT, alkylated with IAA, and digested overnight with trypsin. The resulting peptides were then extracted, dried, and resuspended in 7 μL of 0.1% formic acid. Following centrifugation at 10,000 × g to remove particulates, 5.5 μL of each sample was analyzed using nanoLC-MS/MS (Orbitrap Eclipse). Peptide fragmentation spectra were searched against the human and mouse database using Proteome Discoverer version 2.4 and the Sequest HT search algorithm (Thermo).

### CRISPR/Cas9-mediated genome editing

PRPS1 (P1) and PRPS2 (P2) knockout (KO) clones were created in NIH3T3 cells using a CRISPR/Cas9 nickase system. To clone the crRNAs – BfuAI restriction sites, tracrRNA, and GFP sequences were added into a pLKO.1 - TRC cloning vector. PRPS1 and PRPS2 crRNA oligonucleotides were inserted to a BfuAI-digested pLKO.1 backbone. These plasmids, containing the sgRNA and Cas9 D10A Nickase (Addgene #41816), were co-transfected into NIH3T3 cells. After 48 hours, cells were clonally sorted based on GFP expression onto collagen-coated 96-well plates using BD FACSAria. Clones were assayed for loss of protein expression by Western blotting and were further checked for mutations by purifying genomic DNA and performing PCR on the region spanning the edit site.

For PRPSAP1 (AP1) and PRPSAP2 (AP2) KO clones, NIH3T3 cells were first transduced with a dox-inducible lentiviral Cas9 plasmid (Addgene #50661) and selected with puromycin. Annealed crRNAs were cloned into a lenti-sgRNA vector with hygromycin selection (Addgene #104991) using BsmBI digestion. These lentiviral sgRNA vectors were then introduced into cells expressing dox-inducible Cas9 and selected with hygromycin. Cells were treated with 1 µg/mL doxycycline for a week before clonally sorting them onto collagen-coated 96-well plates using BD FACSAria. Clones were validated for loss of protein expression by Western blotting.

Lentiviral AP2 sgRNA vectors were introduced into validated AP1 KO cell lines to create AP1/AP2 double KO cell lines. P1 and P2 sgRNAs were cloned into a lenti-sgRNA vector with neomycin selection (Addgene #104992) using BsmBI digestion. These sgRNA vectors were then introduced in AP1, AP2 and/or AP1/AP2 KO cell lines to generate the remaining double (AP1/P1, AP1/P2, AP2/P1, AP2/P2) and P2/AP1/AP2 triple KO cell lines. Cells were selected with neomycin and treated with 1 µg/mL doxycycline for a week before clonally sorting them onto collagen-coated 96-well plates using BD FACSAria. Clones were validated for loss of protein expression by Western blotting. All the sgRNAs used are listed in Supplementary Table 6.

CRISPR/Cas9 and a DNA donor repair template were used to generate endogenous PRPS1-mNG11-ALFA knock-in via homology-directed repair (HDR). Plasmid donor DNA was created by inserting two homology arms (∼800 bp each) flanking the exonic mutant sequences into an empty mammalian expression vector (Addgene #68375) using Gibson assembly. PRPS1 crRNA (IDT) was annealed with tracrRNA (IDT) to form sgRNA and incubated with Cas9 protein (IDT) to form a ribonucleoprotein complex (RNP). The RNP mixture, along with the donor plasmid, was electroporated into NIH3T3 cells using the Neon Transfection System (Invitrogen #MPK5000) according to the manufacturer’s instructions. 48 hours after electroporation, cells were transfected with pSFFV_mNG3K (1-10) (Addgene #157993). 36 hours after transfection, correctly knocked-in cells were clonally sorted based on neon green fluorescent protein. ALFA expression was confirmed using immunoblots. For further validation, genotyping was performed by sequencing PCR products amplified around the edit site. All the oligos and primers used are listed in Supplementary Table 6.

### HPA dataset

The Human Protein Atlas (HPA)^72^ tissue consensus database was examined to explore the transcript level of PRPS complex members across various human tissues. The consensus normalized expression levels (nTPM) value for each gene and tissue type represents the maximum nTPM value based on data from both HPA and Genotype-Tissue Expression (GTEx).

### Cell Cycle Analysis

The cells were harvested and washed in ice-cold PBS followed by fixation in 66% ice-cold ethanol solution. Subsequently, the cells were stained with Propidium Iodide (Sigma-Aldrich #P4864) at a final concentration of 50µg/mL and RNase (Roche #111119915001) at a final concentration of 50µg/mL. Samples were then incubated in the dark in a 37°C non-CO2 incubator prior to analysis on BD LSRFortessa. Data was analyzed using FlowJo (BD Biosciences) software with Watson (pragmatic) model.

### Seahorse Assays

Adherent cells were seeded at a density of 10,000 cells/well in DMEM supplemented with 10% FBS and 1X penicillin/streptomycin in a Seahorse XF96 Cell Culture Microplate. The following day, the media was changed to DMEM (Agilent #103575-100) supplemented with 10 mM glucose, 2 mM glutamine, and 1 mM pyruvate and incubated in a 37°C non-CO2 incubator for 1 hour before the assay. Seahorse XF Cell Mito stress Test (Agilent #103015-100) and XF ATP Rate Assay (Agilent #103592-100) were performed according to the manufacturer’s instructions on a Seahorse XFe96 Analyzer (Agilent #S7800B). In both assays, the last injection port was utilized for injecting Hoechst stain. Hoechst fluorescent intensity was measured using a CLARIOstar microplate reader and used to normalize for cell number. Data were exported and analyzed using the Seahorse Wave Desktop Software (Agilent).

### ATP determination assay

Total cellular ATP was measured on a CLARIOstar microplate reader using ATP Determination Kit (Invitrogen #A22066) following manufacturer’s instructions. Protein levels were quantified using BCA reagent (Thermo #23227).

### Soft agar colony formation assay

Cells were seeded at a density of 10,000 cells per well in the top layer of 0.3% agar (Lonza SeaKem LE Agarose #50000) mixed with culture media (Gibco DMEM #12800-058), supplemented with 10% FBS and 1X penicillin/streptomycin, in 6-well plates. The bottom layer consisted of 0.6% agar with the same culture media composition. The top layer was replenished weekly, and images were captured after 3.5 weeks. Image quantification was performed using the Trainable Weka Segmentation Plugin in FIJI^73^. Colonies larger than 9000 µm² were classified as true positives, based on the area of colonies observed in non-transformed NIH3T3 cells. Given the average surface area of an NIH3T3 cell, a cumulative area of 9000 µm² represents at least 35 cells.

### Cell collection and processing for metabolomics

For the stable isotope experiment, NIH-3T3 fibroblasts were seeded in 15 cm plates and cultured in DMEM. The media were replaced with glucose-free media containing 10 mM of ^12^C6-glucose (unlabeled) or ^13^C6-glucose (Cambridge Isotope Laboratories), supplemented with 10% dialyzed FBS and 1X penicillin/streptomycin, and incubated at 37°C for 30 min or 5 hours. After 30 min or 5 hours of isotope exposure, the medium was aspirated, and the cells were washed 3 times with ice-cold PBS. Subsequently, metabolic activity was halted by quenching with ice-cold acetonitrile (CH3CN), followed by the addition of nanopure water (CH3CN: H2O at 2:1.5 (V/V)) to facilitate cell scraping and collection. Polar and non-polar metabolites were extracted using the solvent partition method – acetonitrile: water: chloroform (CH3CN: H2O: CHCl3 at 2:1.5:1 (V/V))^74^. The aqueous phase containing polar metabolites and the organic phase containing non-polar metabolites were separated and dried using vacuum lyophilization (CentriVap Labconco) or a SpeedVac device. The protein pellets were washed in 500 µL of 100% methanol and centrifuged at 12,000 x g at 4°C for 10 min. After discarding the supernatant, the protein pellet was dried in a SpeedVac centrifuge for 20 min. The resulting protein residue pellet was used for normalization.

### NMR data acquisition, processing and analysis

For the analysis of intracellular metabolites, the lyophilized polar extracts were resuspended in 220 μL of NMR buffer containing 100 mM phosphate buffer (pH 7.3), 1 mM trimethylsilyl propionic acid-d4 sodium salt (TSP) as internal standard, and 1 mg/mL sodium azide in 100% deuterium oxide (D2O). 200 μL of each sample was transferred to a 3 mm NMR tube. One-dimensional (1D) ^1^H-NMR spectra were acquired at 288 K on a Bruker Avance III HD 600 MHz spectrometer (Bruker Biospin) equipped with a 5 mm Broad Band Observed (BBO) Prodigy probe. The noesygppr1d pulse sequence was employed with water presaturation (25 Hz bandwidth), 512 transients, a 15-ppm spectral width, a 4.0 s relaxation delay, and a 2.0 s acquisition time. Before Fourier transformation, spectra were zero-filled to 128 K data points and apodized with a 1 Hz exponential line-broadening function. Additionally, 1D ^1^H-^13^C Heteronuclear Single Quantum Correlation (HSQC) spectra were recorded using the hsqcetgppgsisp2.2 pulse sequence with a 15-ppm spectral width, 1024 transients, 1.75 s relaxation delay, and a 0.25 s acquisition time. The spectra were processed with zero-filling to 16 K data points and apodized with unshifted Gaussian function and 4-Hz exponential line broadening. All spectra were recorded and transformed using Topspin 3.6.2 software (Bruker BioSpin, USA) and processed (phased and baseline corrected) using MestReNova software (MNova v12.0.3, Spain). The spectra were internally calibrated to the methyl resonance of the TSP at 0.0 ppm. Metabolites were identified and assigned by comparing with in-house databases, public databases HMDB (Human Metabolome database)^75^, BMRB (Biological Magnetic Resonance Data Bank)^76^, and literature reports. Additionally, 2D ^1^H-^1^H Total Correlation Spectroscopy (TOCSY) experiments were recorded to facilitate the identification of biochemical substances.

To determine the metabolite abundance and their ^13^C isotopomers across the samples, the area of each assigned and well resolved metabolites were manually integrated using global spectra deconvolution (GSD), a line-fitting deconvolution algorithm available in MestReNova software (MNova v12.0.3, Spain), which returns the area of each peak of interest. The peak area was divided by the number of protons contributing to that signal. For absolute quantification the corrected peak areas were converted to molar concentration by calibration against the peak intensity of TSP at 0 ppm for ^1^H spectra and that of lactate methyl group resonance at 1.32 ppm (quantified from 1D ^1^H spectra) for 1D ^1^H-^13^C-HSQC spectra before normalization with milligrams (mg) of protein residue in each sample as a proxy of cell amount.

### Statistics and reproducibility

All statistical analyses were performed using GraphPad Prism software (GraphPad Software). Sample sizes, replicates, and statistical tests (unpaired two-sided Student’s t-test, one-way analysis of variance (ANOVA), and two-way ANOVA) used are indicated in each figure legend. For all statistical analyses, P values are indicated in each figure legend, ns stands for not significant (P > 0.05). Whenever representative western blot images for SEC data are shown, data sets with similar results were generated from at least two independent cell culture/mice tissue harvest experiments in separate SEC runs.

### Data availability

All data supporting the findings of this study are available within the Article and its Supplementary Information, which includes the mass spectrometry proteomics data, the NMR metabolomics data, and the sgRNAs used for generating CRISPR knock-in/knock-out cell lines. For evolutionary analyses, the datasets (sequences; genomic, transcriptomic, or amino acid) generated and/or analyzed during the current study will be available upon request and released during publication.

## Acknowledgements

This work received support by NIH grants (R01CA230904 and R35GM133561) to J.T.C and S10OD026717 to K.D.G. for the mass spectrometer used in the proteomics studies. We thank the Translational Metabolomics Facility at Cincinnati Children’s Hospital Medical Center, and the University of Cincinnati Proteomics laboratory for their assistance. We thank D. Plas, K. Patra, A. Waters, C. Bartolacci, and J. Meller for their insights during the manuscript review process.

The silhouette images in phylogenetic trees were downloaded from Phylopic or designed by Adobe Illustrator software. Schema generation and figure formatting was done using Adobe Illustrator software.

## Author contributions

Conceptualization, B.R.K. and J.T.C; Methodology, B.R.K. and J.T.C.; Investigation, B.R.K., J.T.C., A.C.M., S.V.M.; Formal analysis, B.R.K., A.C.M., S.V.M.; Writing – Original Draft, B. R.K. and J.T.C.; Writing – Review & Editing, B.R.K., J.T.C., A.C.M., S.V.M., K.D.G., L.E.R.; Visualization, B.R.K. and J.T.C.; Project administration, J.T.C.; Supervision, J.T.C., L.E.R., K.D.G.; Funding Acquisition, J.T.C.

## Supplementary Figures and Tables

**Supplementary Fig. 1:**
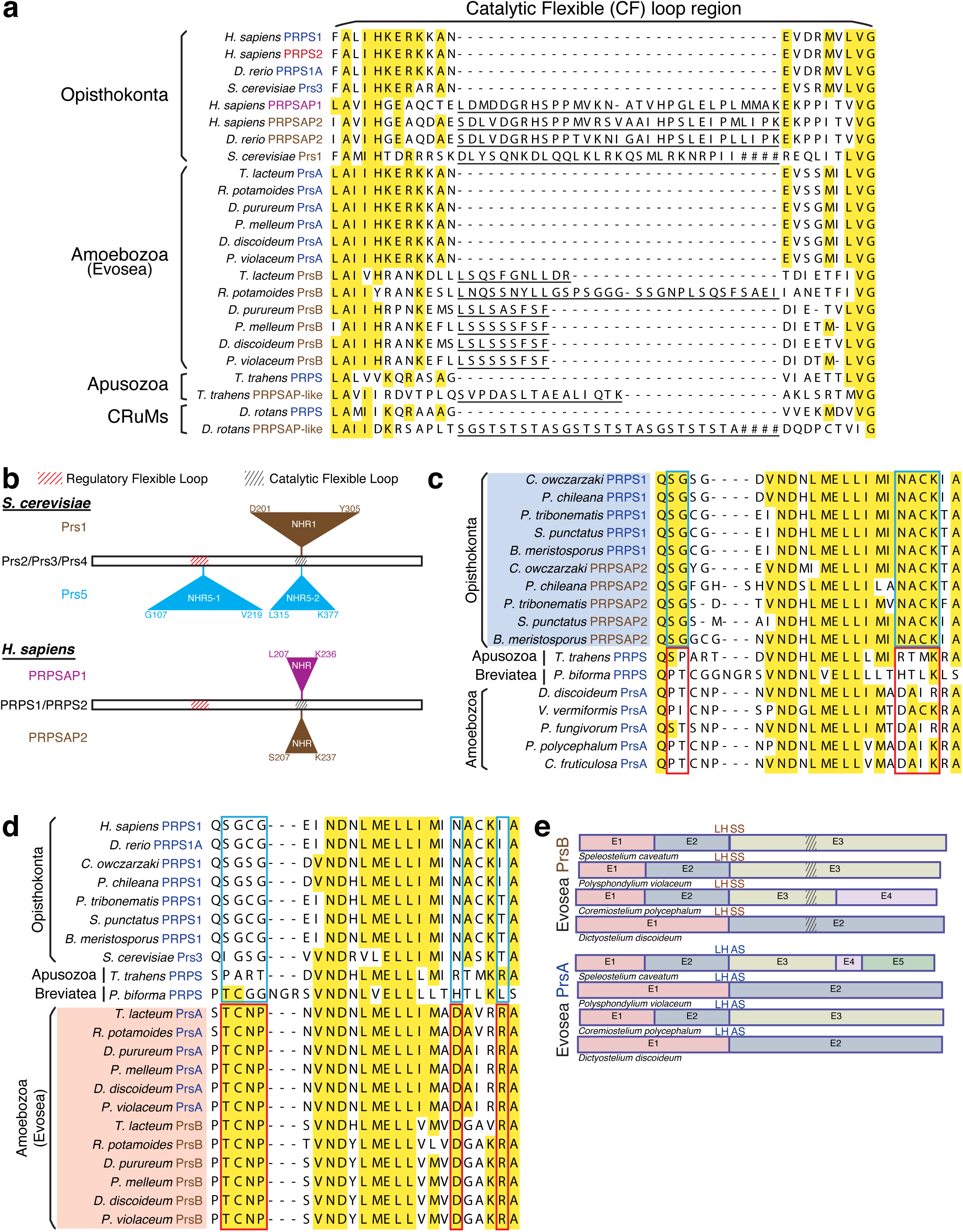
Convergent evolution of PRPSAP-like homologs throughout Amorphea via gene duplications from PRPS1-encoding genes. a) The catalytic flexible (CF) loop from the sequence alignments of PRPS homologs from representative organisms of Opisthokonta, Amoebozoa (Evosea), Apusozoa, and CRuMs. PrsA in Evosea is the PRPS enzyme while PrsB represents the PRPS homologs with insertions in the CF loop (NHRs), similar to Opisthokonta PRPSAP2. PRPS homologs in Apusozoa and CRuMs that share features with Opisthokonta PRPSAP2 are termed “PRPSAP-like”. Underlined residues denote the NHRs. For *S. cerevisiae* Prs1 and *D. rotans* PRPSAP-like sequence, only a part of the insertion is shown, and the rest of the sequence is represented by hatch marks. b) PRPS paralogs in *Saccharomyces cerevisiae* and *Homo sapiens* showing relative positions of non-homologous regions – NHR1 of Prs1, NHR5-1 and NHR5-2 of Prs5, NHRs of PRPSAP1 and PRPSAP2. The open bar represents the polypeptide for Prs2, Prs3, and Prs4 in *S. cerevisiae*, and for PRPS1 and PRPS2 in *H. sapiens*. The insertion points for Prs1, Prs5, PRPSAP1, and PRPSAP2 are marked, with NHRs indicated as triangles either above or below the open bar. The residue numbers on the NHRs indicate the amino acid positions on the respective homologs. c) and d) The N-terminal residues from a sequence alignment of PRPS homologs from representative organisms of Amorphea. Opisthokonta PRPSAP2 sequences are more similar to Opisthokonta PRPS enzymes (blue box) than to PRPS from Apusozoa, Breviatea and Amoebozoa (c) while Evosea PrsB sequences are more similar to Evosea PrsA (red box) than to PRPS from Apusozoa, Breviatea and Opisthokonta (d) indicating that PRPSAP2 and PrsB likely emerged independently from ancestral Opisthokonta PRPS1 and Evosea PrsA, respectively. e) Analysis of conserved splice site junction among PrsA and PrsB homologs across different representative organisms of Evosea. Exons from each of the Prs encoding genes are merged to highlight the splice site junctions. PrsA shares a conserved splice site junction – LH_(A/S)S – with PrsB, further supporting that PrsB likely originated from a gene duplication event in the ancestral Evosea PrsA.

**Supplementary Fig. 2:**
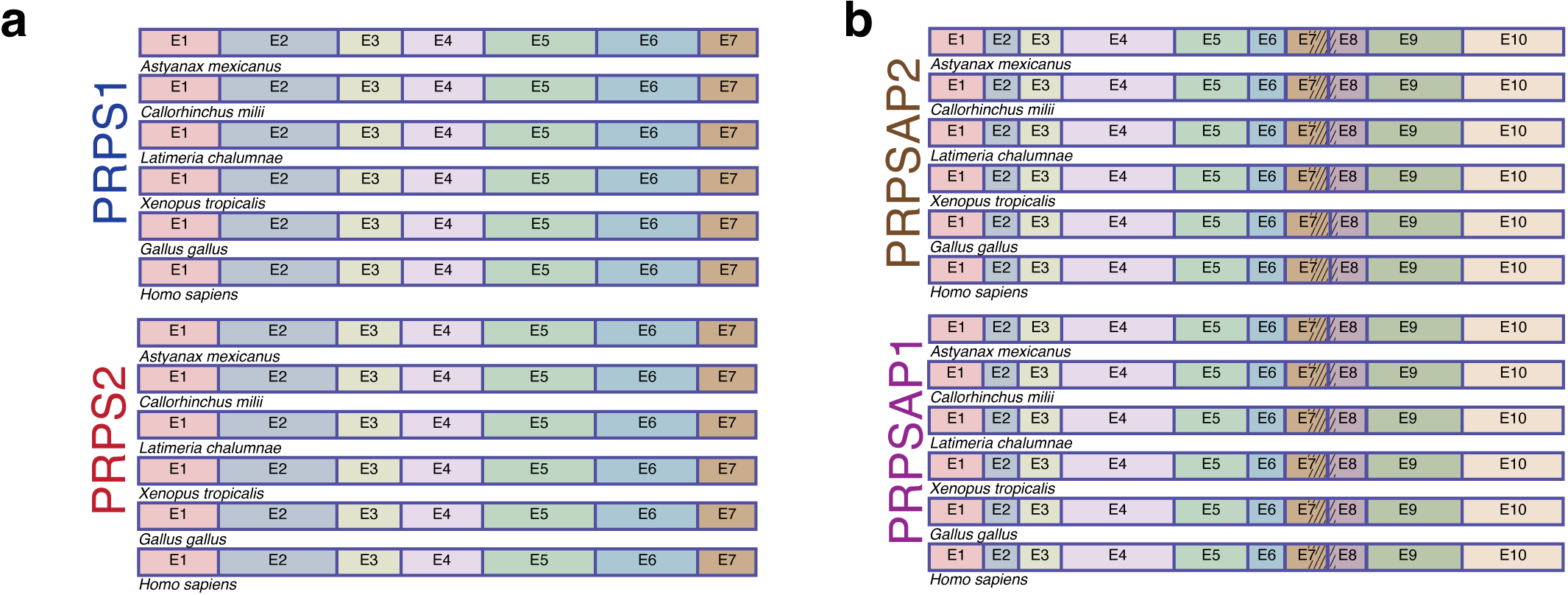
PRPS2 and PRPSAP1 emerged in the ancestor of jawed vertebrates from PRPS1 and PRPSAP2, respectively. a) Analysis of splice site junctions among PRPS1 and PRPS2 across different representative organisms of jawed vertebrates. Exons from each of the PRPS1 and PRPS2 encoding genes are merged to highlight the highly conserved splice site junctions. b) Analysis of splice site junction among PRPSAP2 and PRPSAP1 across different representative organisms of jawed vertebrates. Exons from each of the PRPSAP2 and PRPSAP1 encoding genes are merged to highlight the highly conserved splice site junctions. Hatch marks representing the NHRs are not intended to accurately represent the variable length of the NHRs.

**Supplementary Fig. 3:**
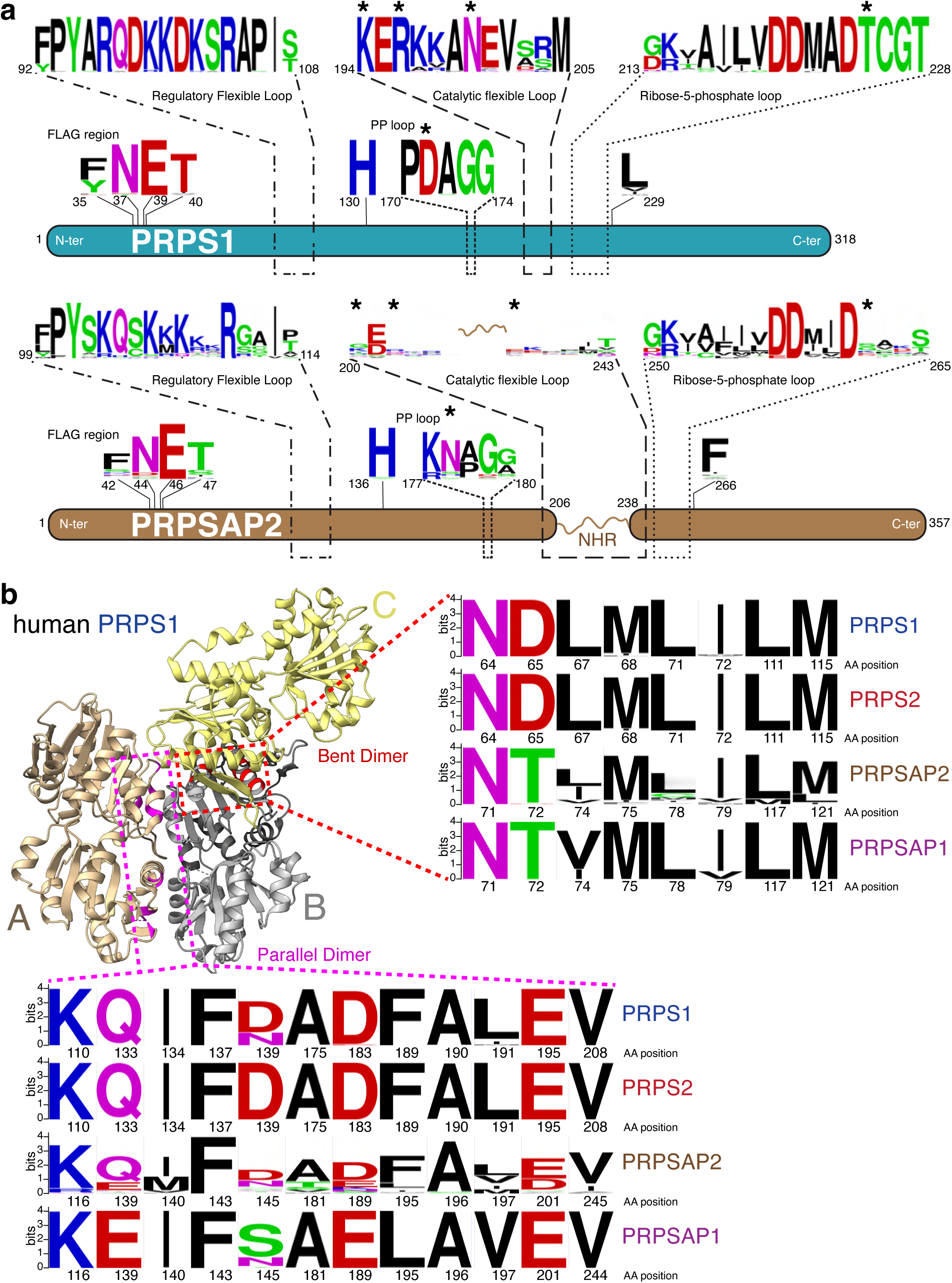
Opisthokont PRPSAP homologs have non-conserved loop regions critical for catalysis but conserved dimer interfaces. a) WebLogo depicting the multiple sequence alignment of the active site residues of PRPS1 and PRPSAP2 from representative organisms in Opisthokonta (n = 44 for PRPS1 and n = 44 for PRPSAP2). Residues from the FLAG region, regulatory flexible loop, pyrophosphate (PP) loop, CF loop, and R5P binding loop are indicated. Many residues shown are conserved in PRPS1 but not in PRPSAP2. The numbers below the logo sequences indicate the corresponding residue positions of the human PRPS1 (NP_002755.1) and PRPSAP2 (NP_001340030.1). Asterisks denote some evolutionarily conserved active site residues that are exclusive to PRPS enzymes and highlighted in Fig.1(d) but replaced with different residues at the corresponding positions in PRPSAPs, highlighted in Fig.1(e). b) The trimeric structure of human PRPS1 (PDB: 2HCR). In the dashed box, red- and magenta-colored residues represent those involved in the formation of bent (B and C subunits) and parallel (A and B subunits) dimers, respectively. The amino acid sequence of *B. subtilis* PRPS^15^ was aligned with the Opisthokonta PRPS homologs, and the corresponding dimer interface residues were selected for generating the WebLogo. Sequences for PRPS1 (n = 44) and PRPSAP2 (n = 44) are derived from representative organisms within opisthokonts, while PRPS2 (n = 46) and PRPSAP1 (n = 92) sequences are derived from representative organisms within jawed vertebrates. The numbers below the logo sequences indicate the corresponding residues positions of the human PRPS1 (NP_002755.1), PRPS2 (NP_002756.1), PRPSAP1 (AAH09012.1), and PRPSAP2 (NP_001340030.1). The significant sequence conservation between PRPS and PRPSAP indicates the potential for heteromeric associations.

**Supplementary Fig. 4:**
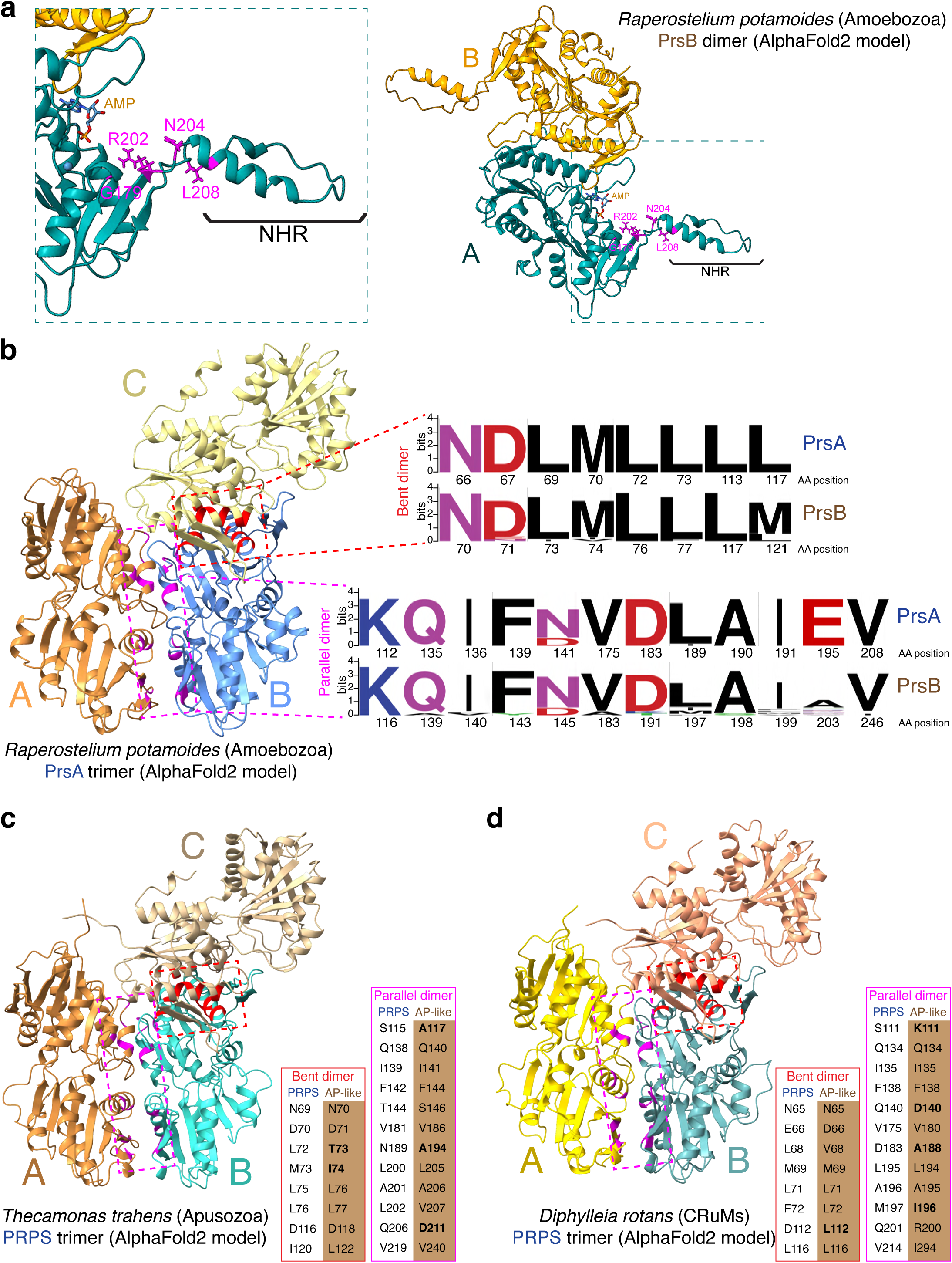
PRPSAP-like homologs in non-opisthokont Amorphea lineages are likely non-catalytic with conserved dimer interfaces. a) Predicted dimeric structure of *R. potamoides* PrsB (annotated from SRX8374346-9) from the AlphaFold2 model, and a zoom in highlights four non-conserved residues in PrsB (magenta) at the corresponding positions of residues for PrsA (Table 1). AMP was modeled into the dimer to denote the putative ATP binding site. NHR represents the insertion in the CF loop. b) Predicted trimeric structure of *R. potamoides* PrsA (annotated from SRX8374346-9) from the AlphaFold2 model. In the dashed box, red- and magenta-colored residues represent those involved in the formation of bent (B and C subunits) and parallel (A and B subunits) dimers, respectively. The amino acid sequence of *B. subtilis* PRPS^15^ was aligned with the Amoebozoa PRPS homologs, and the corresponding dimer interface residues were selected for generating the WebLogo. Sequences for PrsA (n = 19) and PrsB (n = 20) are derived from representative organisms within Amoebozoa. The numbers below the logo sequences indicate the corresponding residues positions of *R. potamoides* PrsA and PrsB. Significant sequence conservation between PrsA and PrsB indicates the potential for heteromeric associations. c) and d) represents the predicted trimeric structure of *T. trahens* (representative organism of Apusozoa) PRPS (XP_013753676.1) in (c) and *D. rotans* (representative organism of CRuMs) PRPS (annotated from SRX3153023) in (d) from the AlphaFold2 model. In the dashed box, red- and magenta-colored residues represent those involved in the formation of bent (B and C subunits) and parallel (A and B subunits) dimers, respectively. A comparison of dimer interface residues at similar positions in PRPSAP-like is shown. Non-conserved residues in PRPSAP-like relative to PRPS are shown in bold. Significant sequence conservation between PRPS and PRPSAP-like indicates the potential for heteromeric associations.

**Supplementary Fig. 5:**
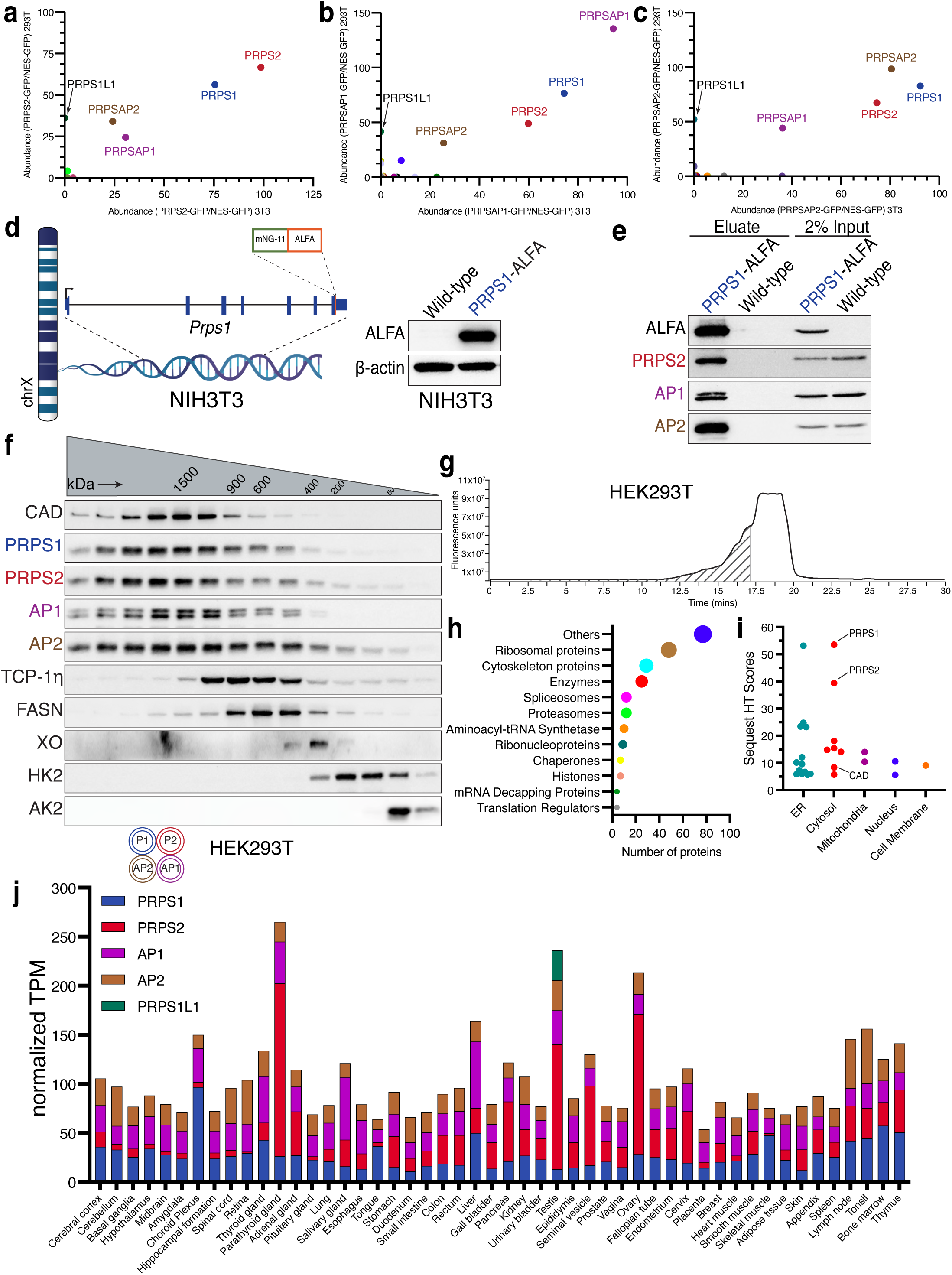
The stable PRPS complex is one of the largest cytosolic metabolic assemblies in cells. a) and b) and c) represent the scatter plots from mass spectrometry (MS) runs of eluates from GFP IP in stably expressing PRPS2-GFP, PRPSAP1-GFP, and PRPSAP2-GFP, respectively in NIH3T3 (x-axis) and HEK293T (y-axis) cells. The axes represent the square root-transformed SEQUEST HT scores normalized to control. d) Schematic of *Prps1* gene with an endogenous ALFA tag knocked in frame at the C-terminus. mNG-11 represents the short fragment of the split monomeric Neon Green protein and ALFA is an epitope tag containing residues – PSRLEEELRRRLTEP. Western blot validating full-length expression of the endogenously tagged PRPS1-ALFA protein is shown. e) ALFA pulldown from the whole cell extracts of knock-in NIH3T3 cells demonstrating the interaction of endogenous PRPS1 with PRPS2, PRPSAP1, and PRPSAP2. f) Western blot analysis of SEC fractions collected from HEK293T native whole cell lysates. Cell lysates were fractionated on a Superose 6 Increase 3.2/300 column. Immunoblots probing PRPS complex members and internal standards are shown. In the pictogram, a double circle means multiple copies of the protein are interacting within the heteromeric complex. g) Chromatogram showing SEC traces of HEK293T whole cell lysates fractionated on a Bio SEC-5 2000Å column, which offers better fractionation for proteins/protein complexes in high molecular weight (HMW) range. The proteins that eluted in the fractions shown as hatch marks were sent for mass-spectrometry analyses for identification of proteins in HMW range. h) Classification of HMW proteins based on their functions (manual curation) from the mass-spectrometry dataset obtained from fractions collected in (g). i) Subclassification of enzymes from (h) based on their cellular localization. j) Normalized transcript per million (nTPM) of PRPS complex components in various human tissues obtained from Human Protein Atlas.

**Supplementary Fig. 6:**
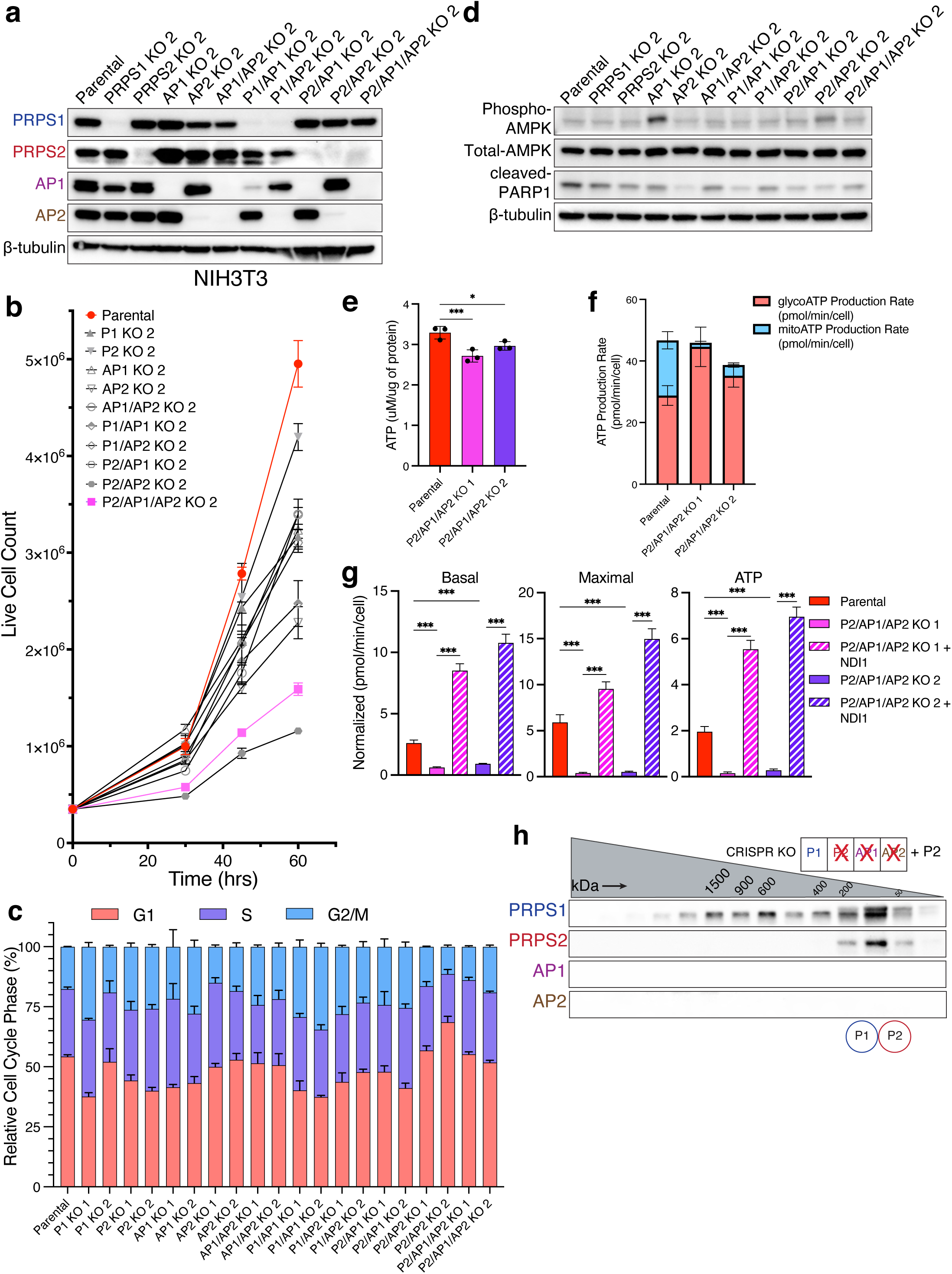
Additional phenotypic characterization of NIH3T3 isogenic knockout lines. a) Western blot validation of the second set of CRISPR-Cas9-generated isogenic knockout cell lines. b) Proliferation for the panel of NIH3T3 knockout cell lines generated in a). Error bars represent SD; n = 3. c) Bar graph depicting propidium-iodide-based cell cycle profiles: G1, S, and G2/M phases for the panel of NIH3T3 knockout cell lines. Error bars represent SD; n = 3. d) Immunoblots were probed with phospho-AMPK (T172) and cleaved PARP1 antibody in the panel of NIH3T3 knockout cell lines as readouts for energy stress and apoptosis, respectively. e) Total cellular ATP (normalized to the protein content) measured by ATP determination assay in NIH3T3 parental and P2/AP1/AP2 KO cell lines. Error bars represent SD; n = 3. (*P < 0.05; ***P < 0.001 by one-way ANOVA post hoc test). f) ATP production rate determined by Seahorse ATP Rate assay in NIH3T3 parental and P2/AP1/AP2 KO cell lines. Error bars represent SD; n = 7. g) Quantification of basal respiration, maximal respiration (post FCCP injection), and respiration coupled to ATP production measured by Seahorse ATP Rate Assay in NDI1 expressing P2/AP1/AP2 KO cell lines. Error bars represent SD; n = 7. (***P < 0.001 by one-way ANOVA post hoc test). h) Western blot analysis of SEC fractions collected from NIH3T3 P2/AP1/AP2 KO native whole cell lysates transiently transfected with PRPS2. Cell lysates were fractionated on a Superose 6 Increase 3.2/300 column. In the pictogram, a single circle means a single protein is interacting within the complex.

**Supplementary Fig. 7:**
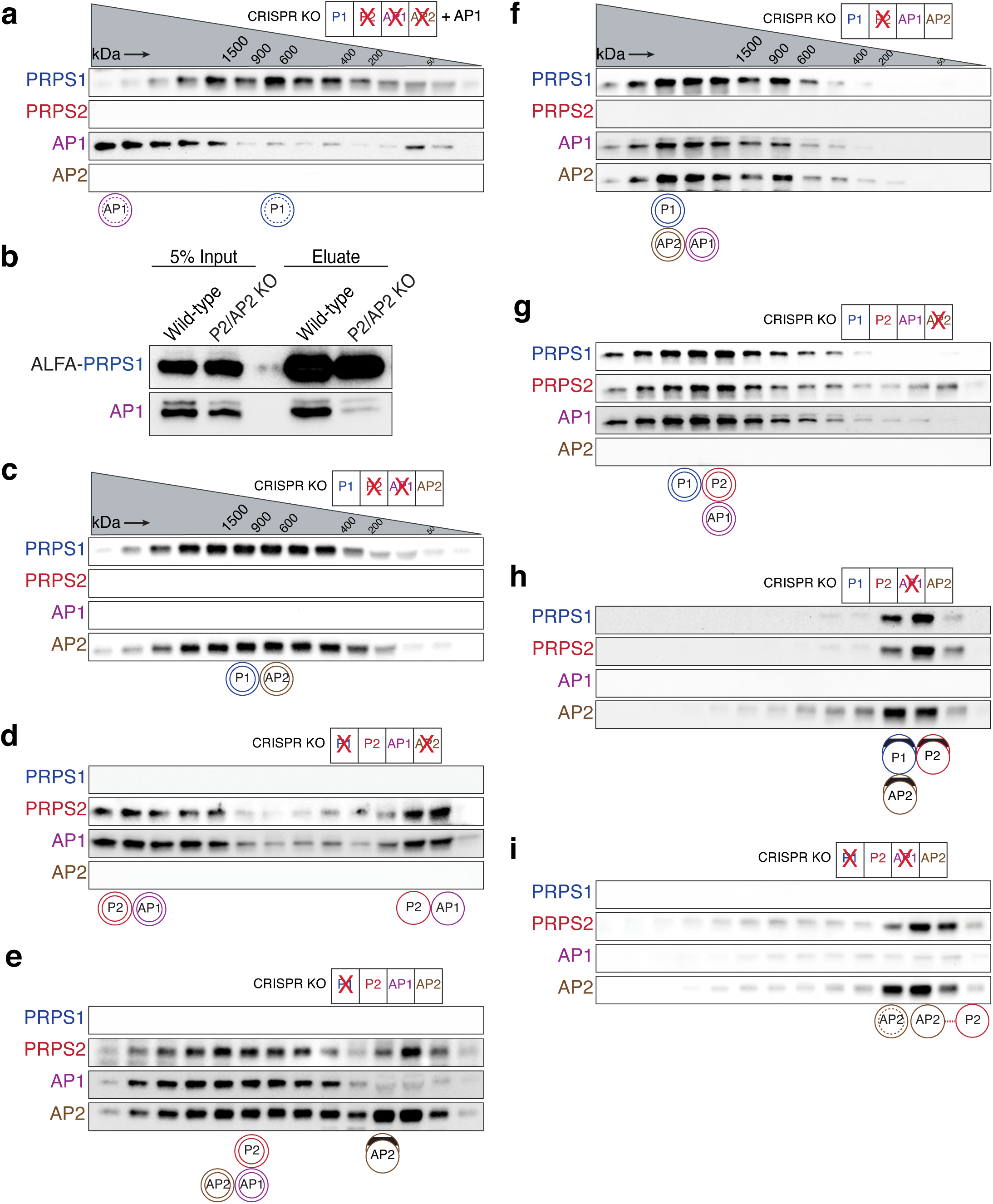
SEC profiles of additional isogenic knockout lines. a) Western blot analysis of SEC fractions collected from native whole cell lysates of NIH3T3 P2/AP1/AP2 KO cells stably expressing AP1 b) ALFA pulldown from the whole cell extracts of NIH3T3 parental and P2/AP2 KO cells transiently transfected with PRPS1-ALFA c-i) Western blot analysis of SEC fractions collected from native whole cell lysates of NIH3T3 P2/AP1 KO cells (c), P1/AP2 KO cells (d), P1 KO cells (e), P2 KO cells (f), AP2 KO cells (g), AP1 KO cells (h), and P1/AP1 KO cells (i). All the fractionation experiments were performed on Superose 6 Increase 3.2/300 column. The circular pictograms at the bottom of the SEC immunoblots illustrate the different configurations of the PRPS complex. A double circle means multiple copies of the protein are interacting within the heteromeric complex. A double circle with a dotted inner circle means multiple copies of the protein are forming homo-oligomers. A single circle means a single protein is interacting within the complex. A circle with lines inside indicates that the proteins may be forming a trimer or tetramer.

**Supplementary Fig. 8:**
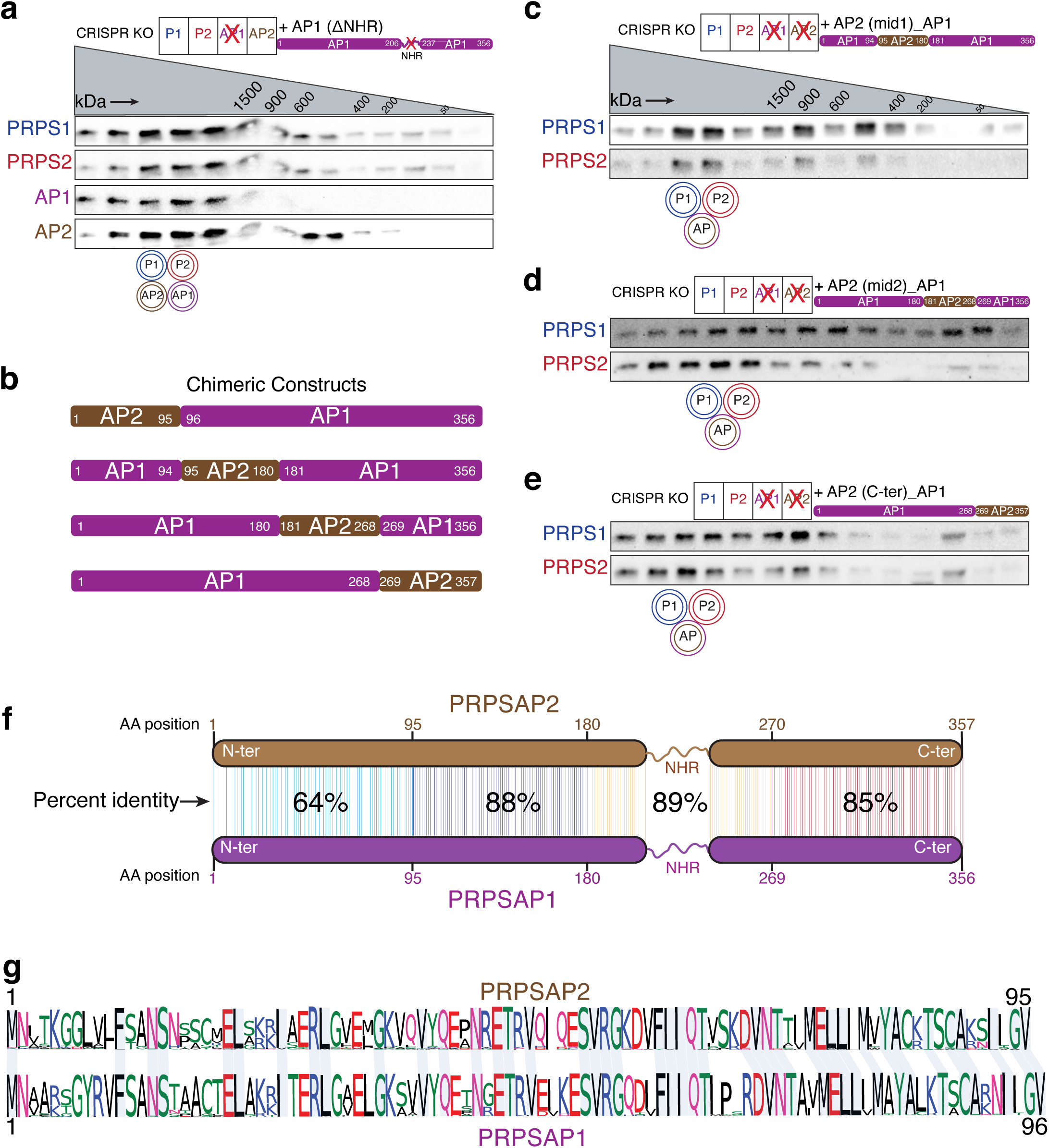
SEC profiles of chimeric PRPSAP1/AP2-expressing cells. a) Western blot analysis of SEC fractions collected from native whole-cell lysates of NIH3T3 AP1 KO cells stably expressing AP1 lacking the non-homologous region (NHR). The absence of NHR in AP1 does not prevent AP1-mediated complex elongation. b) Schema of chimeric constructs created to test AP1-specific domains that confer complex elongation property. c-e) Western blot analysis of SEC fractions collected from native whole-cell lysates of NIH3T3 AP1/AP2 KO cells stably expressing chimeric AP1 constructs: one containing residues 95-180 from AP2 (b), residues 181-268 from AP2 (c), and AP2’s C-terminus (residues 269-357) (d). f) Comparison of human AP1 and AP2 amino acid sequence. The four regions switched in Fig.4 (e) and Supplementary Figs. 8 (c)-(e) are highlighted with different colors. Each line connecting AP1 and AP2 represents identical amino acids at that position. The regions spanning residues 1-95 show the greatest variation, with only 64% amino acid identity. The poorly conserved, highly variable amino acids present in the NHRs were excluded from this analysis. g) MetaLogo depicting the multiple sequence alignment of the N-terminal amino acid residues of AP1 and AP2 from representative organisms of jawed vertebrates (n = 92 for AP1 and n = 93 for AP2). Residue numbers for AP1 and AP2 correspond to the human homologs (AAH09012.1 and NP_001340030.1, respectively). All the fractionation experiments were performed on Superose 6 Increase 3.2/300 column. In the pictogram, a double circle means multiple copies of the protein are interacting within the heteromeric complex.

**Supplementary Fig. 9:**
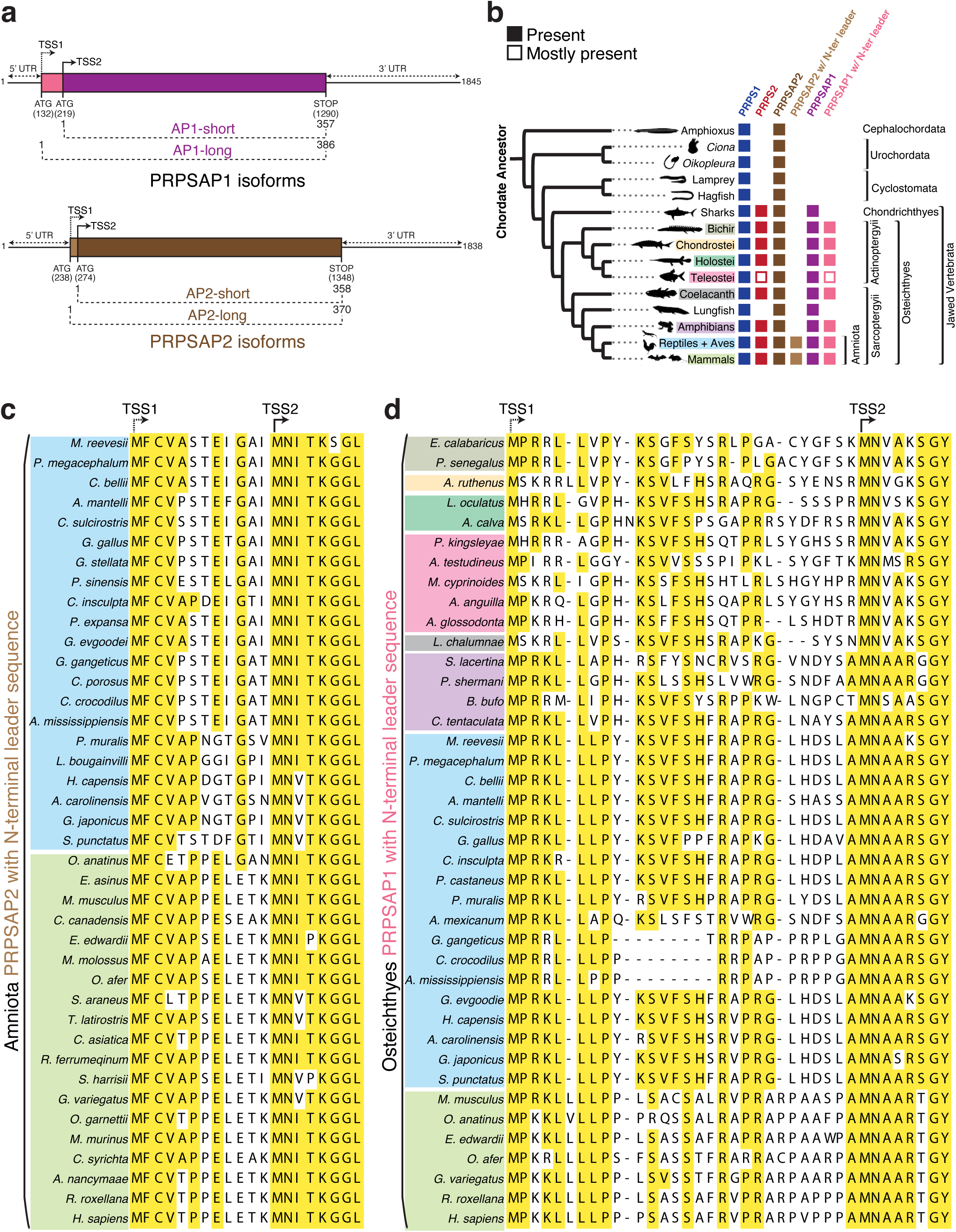
Conservation of upstream translation start site in PRPSAPs. a) Schematic representation of alternative start sites in mammalian AP1 and AP2 and their consequent translation into the short and long isoforms. Base positions for AP1 and AP2 correspond to the mouse homologs. TSS1 and TSS2 represents the upstream and downstream translation start sites, respectively. b) Phylogenetic distribution profiles of PRPS homologs (PRPS1, PRPS2, PRPSAP2, PRPSAP2 with N-terminal leader sequence, PRPSAP1, and PRPSAP1 with N-terminal leader sequence) in chordates (presence/absence) are noted across the tree. PRPSAP2 and PRPSAP1 isoforms with additional N-terminal leader sequences emerged in the ancestor of Amniota and Osteichthyes, respectively. c) and d) The N-terminal residues from a sequence alignment of PRPSAP2 (c) and PRPSAP1 (d) from representative organisms of Amniota and Osteichthyes, respectively. TSS1 and TSS2 represents the upstream and downstream translation start sites, respectively.

**Supplementary Fig. 10:**
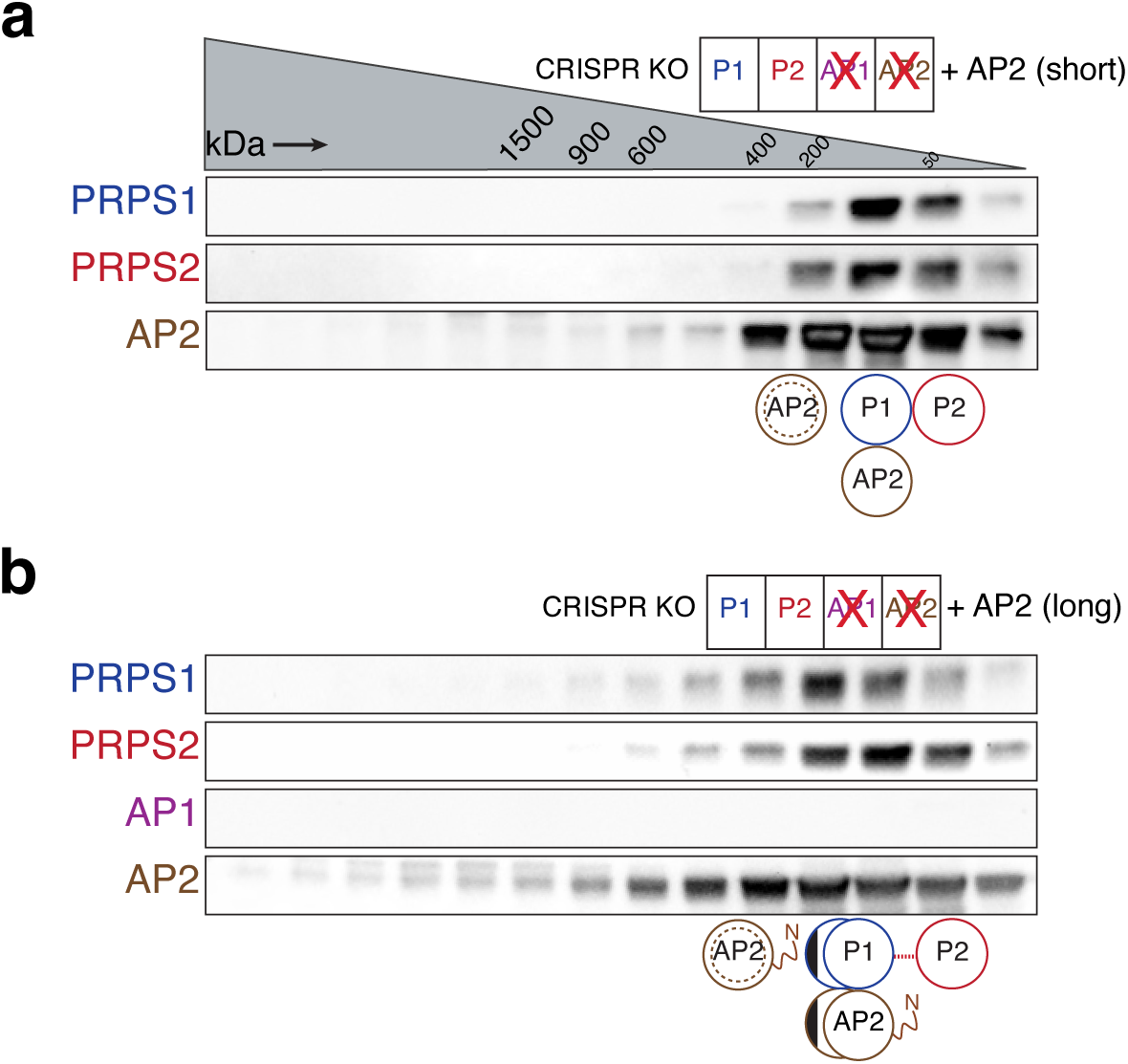
SEC profiles of long and short PRPSAP2 isoforms. a) and b) Western blot analysis of SEC fractions collected from NIH3T3 AP1/AP2 KO cells stably expressing the short isoform of AP2 (a) and long isoform of AP2 (b). Cell lysates were fractionated on a Superose 6 Increase 3.2/300 column. The circular pictograms at the bottom of the SEC immunoblots illustrate the different configurations of the PRPS complex. A double circle with a dotted inner circle means multiple copies of the protein are forming homo-oligomers. A single circle means a single protein is interacting within the complex. A circle with lines inside indicates that the proteins may be forming a trimer or tetramer.

**Supplementary Fig. 11:**
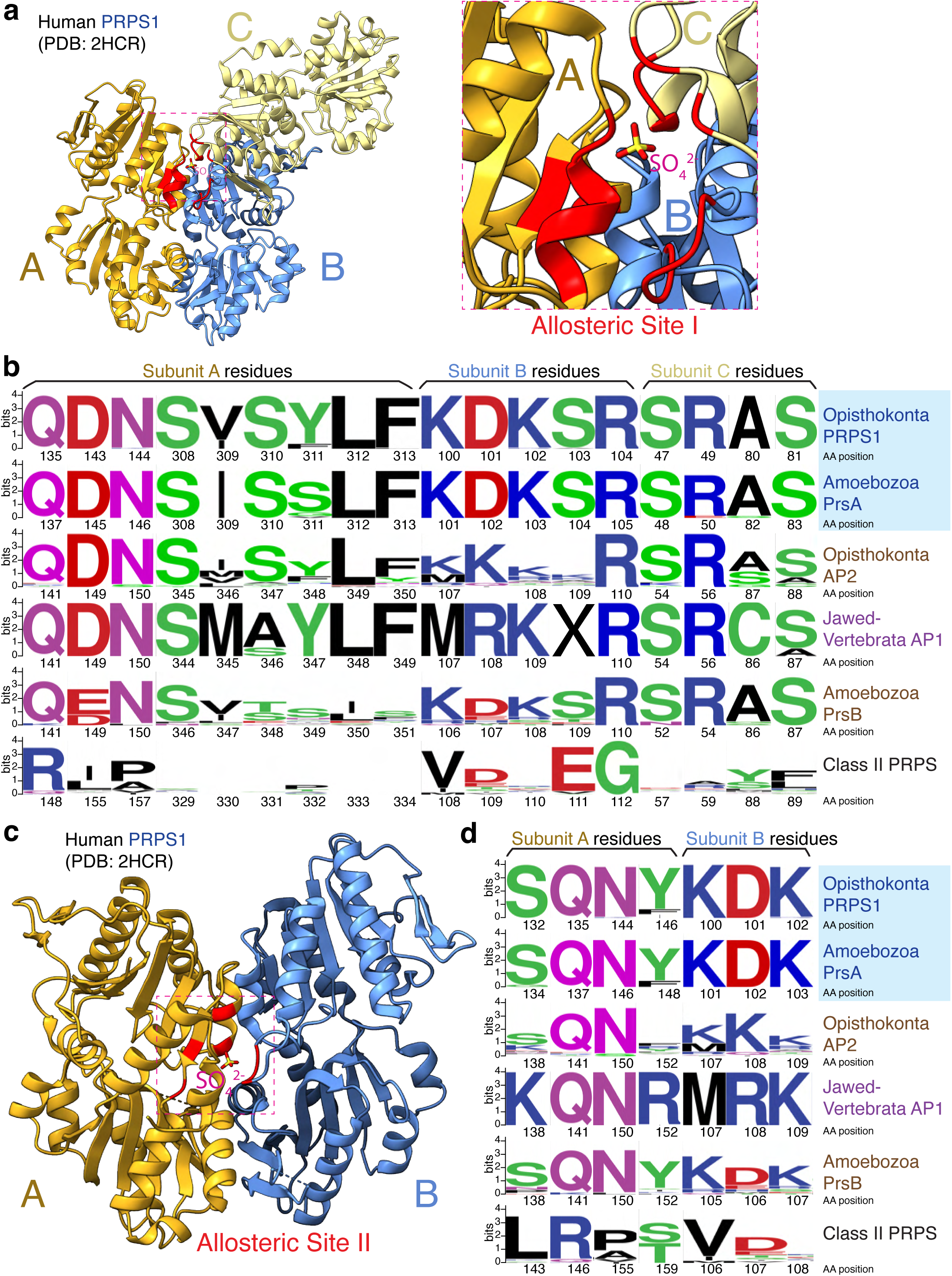
Comparison of allosteric site across PRPS homologs. a) The structure of trimeric human PRPS1 (PDB ID: 2HCR). The dashed box represents the allosteric site I, a zoom in of this site shows SO ^2^ (which represents the phosphate of ADP) positioned at the trimeric interface and red color indicates the residues from each subunit contributing to the formation of allosteric site I. b) The amino acid sequence of the human PRPS1 was aligned with sequences of representative organisms from Opisthokonta PRPS1 (n = 44), Amoebozoa PrsA (n = 19), Opisthokonta PRPSAP2 (n = 46), jawed Vertebrata PRPSAP1 (n = 92), Amoebozoa PrsB (n = 20) and Class II PRPS (n = 53), and the corresponding allosteric site I residues were selected for generating the WebLogo. The numbers below the logo sequences indicate the corresponding residues positions of human PRPS1 and PRPSAP2 (for Opisthokonts), *R. potamoides* PrsA and PrsB (for Amoebozoa), human PRPSAP1 (for jawed Vertebrata), and *A. parasiticum* Class II PRPS annotated from SRX179384. X represents the absent amino acid residue at that position jawed Vertebrata AP1. Class II PRPS enzymes are shown as a reference since they are known to lack allosteric sites found in Class I PRPS (“classical” PRPS enzymes)^2^. c) The structure of dimeric human PRPS1 (PDB ID: 2HCR). The dashed box represents the allosteric site II, a zoom in of this site shows SO ^2^ positioned at the dimer interface and red color indicates the residues from each subunit contributing to the formation of allosteric site II. d) The amino acid sequence of the human PRPS1 was aligned with sequences from Opisthokonta PRPS1, Amoebozoa PrsA, Opisthokonta PRPSAP2, jawed Vertebrata PRPSAP1, Amoebozoa PrsB and Class II PRPS, and the corresponding allosteric site II residues were selected for generating the WebLogo similar to (b).

**Supplementary Fig. 12:**
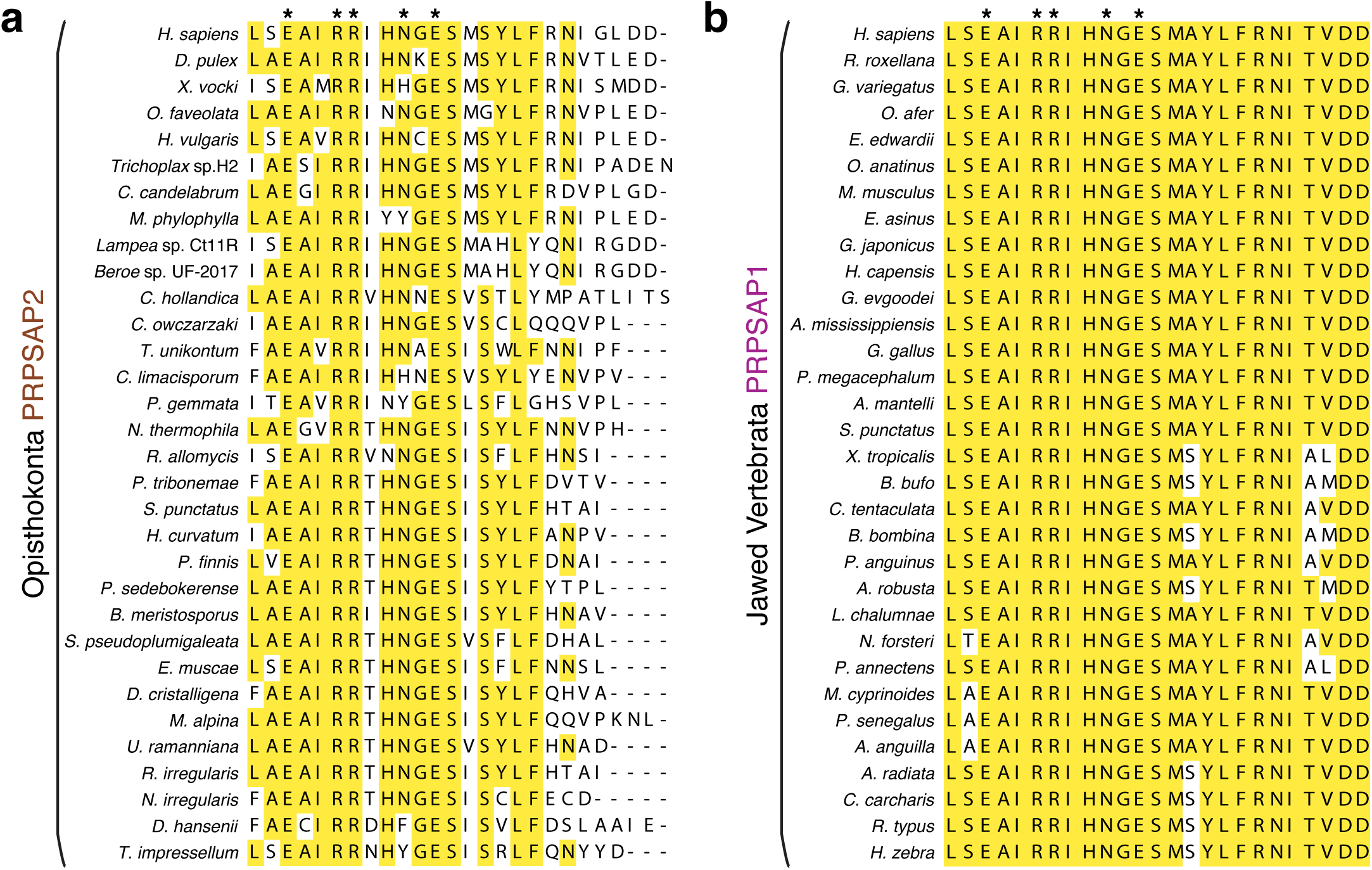
Conservation of filament interfaces in PRPSAPs. a) and b) The C-terminal residues from the sequence alignments of representative organisms from Opisthokonta PRPSAP2 (a) and jawed Vertebrata PRPSAP1 (b). Residues, marked with an asterisk, critical for hexamer stacking at the filament interface are deeply conserved in PRPSAPs.

**Supplementary Fig.13:**
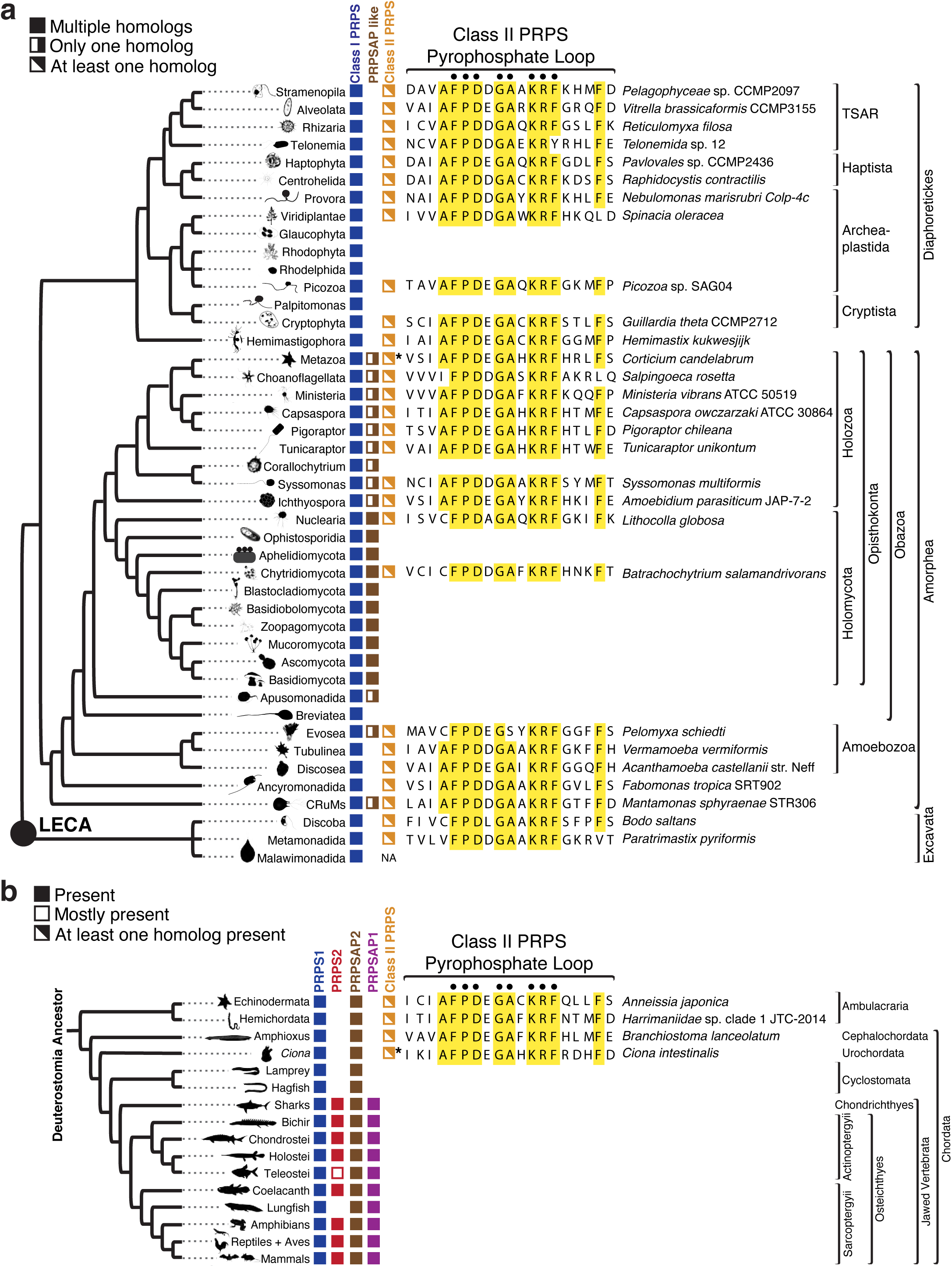
Loss of Class II PRPS enzyme associates with expanded Class I PRPS homolog repertoire. a) Eukaryotic phylogenetic tree of members of Class I PRPS and PRPAP-like as well as Class II PRPS. The presence of single or multiple homologs, that we have detected so far, are noted across the tree. PRPSAP-like orthologs are categorized based on non-conserved active site residues and insertions of varied length in the CF loop. Asterisk in Metazoan Class II PRPS enzyme indicates that it is present in most metazoans except Craniata. Majority of Holomycota species lost Class II PRPS, that do not possess allosteric sites, while expanding multiple Class I PRPS homologs. Sequence alignment of residues surrounding pyrophosphate (PP) loop of class II PRPS for representative organisms of each clade are shown. Conserved residues indicated by solid dots are unique to Class II PRPS. b) Phylogenetic distribution profiles of Class I (PRPS1, PRPS2, PRPSAP1, PRPSAP2) and Class II in deuterostomes (presence/absence) are noted across the tree. In the Urochordata class, Class II PRPS is generally found in most organisms, except for *Oikopleura* which interestingly possess additional Class I PRPS homologs compared to other organisms in this clade. Sequence alignment of residues surrounding pyrophosphate (PP) loop of class II PRPS for representative organisms of each clade is shown. Conserved residues indicated by solid dots are unique to Class II PRPS enzymes. Following the loss of Class II PRPS in Cyclostomata, PRPS2 and PRPSAP1 emerged from gene duplications events in the ancestor of jawed vertebrates.

**Supplementary Table.1:**
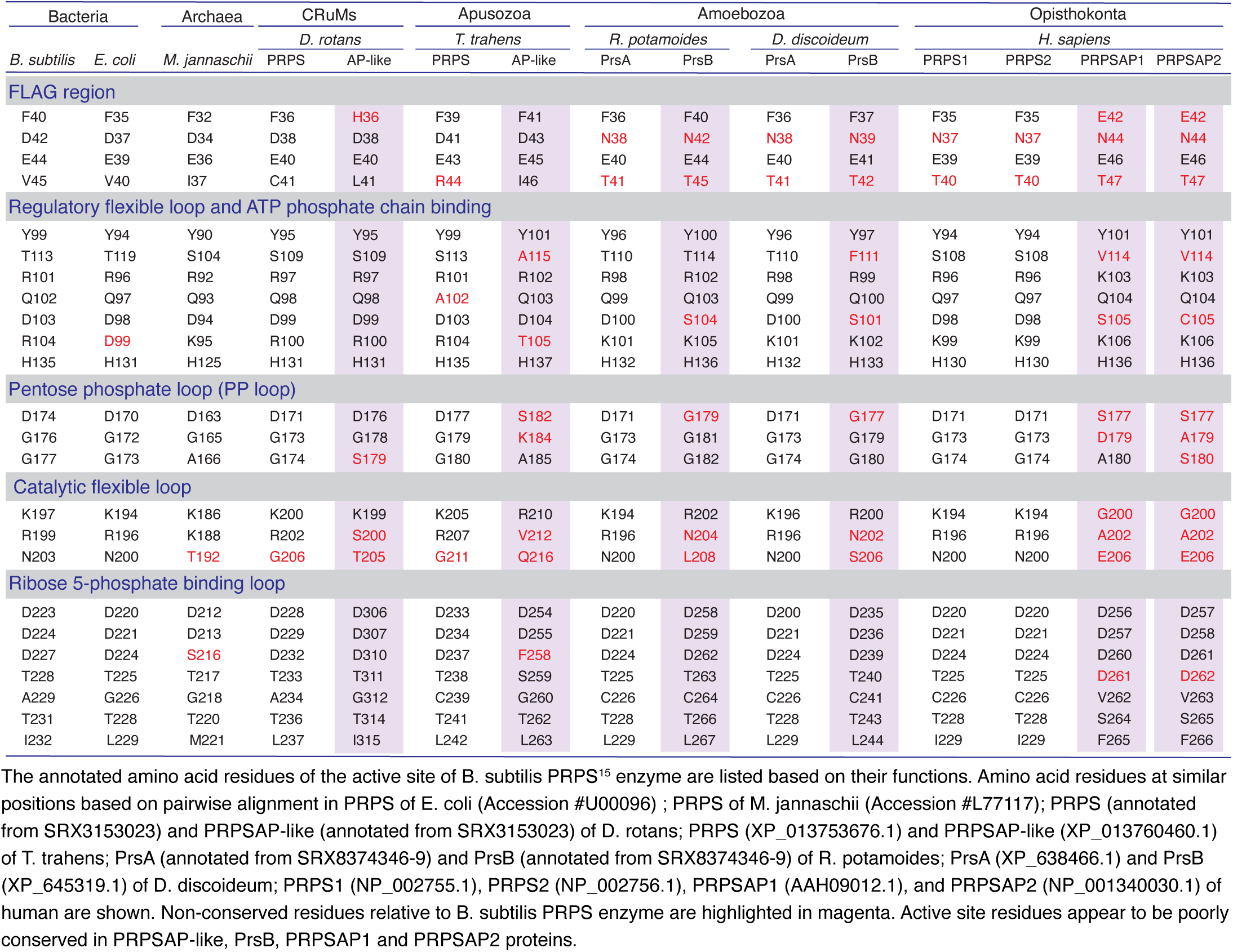
Comparion of active site residues of *B. subtilis* PRPS with corresponding amino acid residues of other Class I PRPS homolgs.

